# Fitness effects of altering gene expression noise in *Saccharomyces cerevisiae*

**DOI:** 10.1101/294603

**Authors:** Fabien Duveau, Andrea Hodgins-Davis, Brian P.H. Metzger, Bing Yang, Stephen Tryban, Elizabeth A. Walker, Patricia Lybrook, Patricia J. Wittkopp

## Abstract

Gene expression noise is an evolvable property of biological systems that describes differences in gene expression among genetically identical cells in the same environment. Prior work has shown that expression noise is heritable and can be shaped by natural selection, but the impact of variation in expression noise on organismal fitness has proven difficult to measure. Here, we quantify the fitness effects of altering expression noise for the *TDH3* gene in *Saccharomyces cerevisiae*. We show that increases in expression noise can be deleterious or beneficial depending on the difference between the average expression level of a genotype and the expression level maximizing fitness. We also show that a simple model relating single-cell expression levels to population growth produces patterns that are consistent with our empirical data. We use this model to explore a broad range of average expression levels and expression noise, providing additional insight into the fitness effects of variation in expression noise.

## Introduction

Gene expression is a dynamic process that results from a succession of stochastic biochemical events, including availability of transcription factors, binding of transcription factors to promoter sequences, recruitment of transcriptional machinery, transcriptional elongation, mRNA degradation, protein synthesis, and proteolysis. These events cause the expression level of a gene product to differ even among genetically identical cells grown in the same environment (Elowitz *et al.* 2002; Chong *et al.* 2015). This variability in gene expression is known as ‘expression noise’ and is under genetic control (Raser and O’Shea 2004; Sanchez and Golding 2013), with heritable variation causing differences in noise among genes (Newman *et al.* 2006) and genotypes (Murphy *et al.* 2007; Hornung *et al.* 2012; Fehrmann *et al.* 2013; Sharon *et al.* 2014; Liu *et al.*2015).

Because gene expression noise is heritable and variable within populations, it can evolve in response to natural selection if it affects fitness. Indeed, prior studies have suggested that expression noise can be either beneficial or deleterious depending on the context (reviewed in Viney and Reece 2013; Richard and Yvert 2014; Liu *et al.* 2016). For example, Metzger *et al.* (2015) provides evidence that increased expression noise can be selected against in natural populations, presumably because elevated noise increases the probability that a given cell produces a suboptimal level of protein expression (Wang and Zhang 2011; Duveau *et al.* 2017). Consistent with this hypothesis, a negative correlation exists at the genomic scale between the expression noise of genes and their dosage sensitivity (Fraser *et al.* 2004; Batada and Hurst 2007; Lehner 2008; Keren *et al.* 2016). However, because the optimal level of gene expression can vary among environments, high gene expression noise has been suggested to be beneficial if it can produce individuals with phenotypes that are better adapted to the environment than individuals produced with low gene expression noise. For instance, noise in gene expression can allow a small fraction of cells to survive when confronted with stressful environmental conditions (Blake *et al.* 2006; Fraser and Kærn 2009; Ito *et al.* 2009; Levy *et al.* 2012; Viney and Reece 2013; Liu *et al.* 2015; Wolf *et al.* 2015). Consistent with this idea, a genomic screen in yeast found that plasma-membrane transporters involved in cell-environment interactions displayed elevated expression noise in yeast (Zhang *et al.*2009). Theoretical work also suggests the existence of cost-benefit tradeoffs that can make expression noise either beneficial or deleterious under different circumstances (Tǎnase-Nicola and Ten Wolde 2008).

Despite a growing body of evidence that selection has acted on expression noise for many genes, direct measurements of how changes in expression noise impact fitness remain scarce (Liu *et al.* 2016). A major reason for this scarcity is that most mutations that alter gene expression noise also alter average expression level (Newman *et al.* 2006; Hornung *et al.* 2012; Carey *et al.* 2013; Sharon *et al.* 2014), making it difficult to disentangle the fitness effects of changing expression noise and average expression level. Here, we directly estimate the effects of changing expression noise on fitness independently from changes in average expression level for the *TDH3* gene of *Saccharomyces cerevisiae*. *TDH3* encodes an isozyme of the yeast glyceraldehyde-3-phosphate dehydrogenase (GAPDH) involved in glycolysis and gluconeogenesis (McAlister and Holland 1985) as well as transcriptional silencing (Ringel *et al.* 2013), RNA-binding (Shen *et al.* 2014) and possibly antimicrobial defense (Branco *et al.* 2014). Variation in this gene’s promoter affecting expression noise has previously been shown to be a target of selection in natural populations (Metzger *et al.* 2015). To assess the impact of changes in expression noise on fitness at different expression levels, we generated mutant alleles of the *TDH3* promoter that covered a broad range of average expression levels and expression noise. We find that increases in expression noise are detrimental when the average expression level of a genotype is close to the fitness optimum, but beneficial when the average expression level of a genotype is further from this optimum. This pattern was reproduced by a simple computational model that links expression in single cells to their doubling time to predict population fitness. We used this individual-based model to explore the fitness effects of a broader combination of average expression levels and expression noise than were explored empirically, showing that not only do the fitness effects of changing expression noise depend on the average expression level, but that the fitness effects of changing average expression level also depend upon the amount of expression noise.

## Results and Discussion

### Generating variation in expression noise independent of average expression level

To disentangle the effects of changes in average expression level and expression noise on fitness, we examined a set of *TDH3* promoter (*P*_*TDH3*_) alleles with a broad range of activities. For each allele, we measured the average expression level and expression noise by cloning the allele upstream of a yellow fluorescent protein (YFP) coding sequence, integrating this reporter gene (*P*_*TDH3*_-*YFP*) into the *HO* locus, and quantifying fluorescence in living cells using flow cytometry in six replicate populations per genotype (Figure 1A). The fluorescence value of each cell was transformed into an estimated mRNA level (Figure 1A) based on the relationship between fluorescence and YFP mRNA abundance (Figure 1B,C). The average expression level of a genotype was then calculated by averaging the median values from the six replicates (Figure 1A) and expressing this value as a percent change from the wild type allele. Expression noise was calculated for each replicate as the variance divided by the median expression among cells, a measure of noise strength similar to the Fano factor (Sanchez and Golding 2013). The expression noise of each genotype was then calculated by averaging the noise strength from the six replicate populations, and this value was expressed as a percent change from the wild type allele. The main conclusions of this study are robust to the choice of noise metric, as shown in supplementary figures using three alternative metrics of noise.

**Figure 1.**
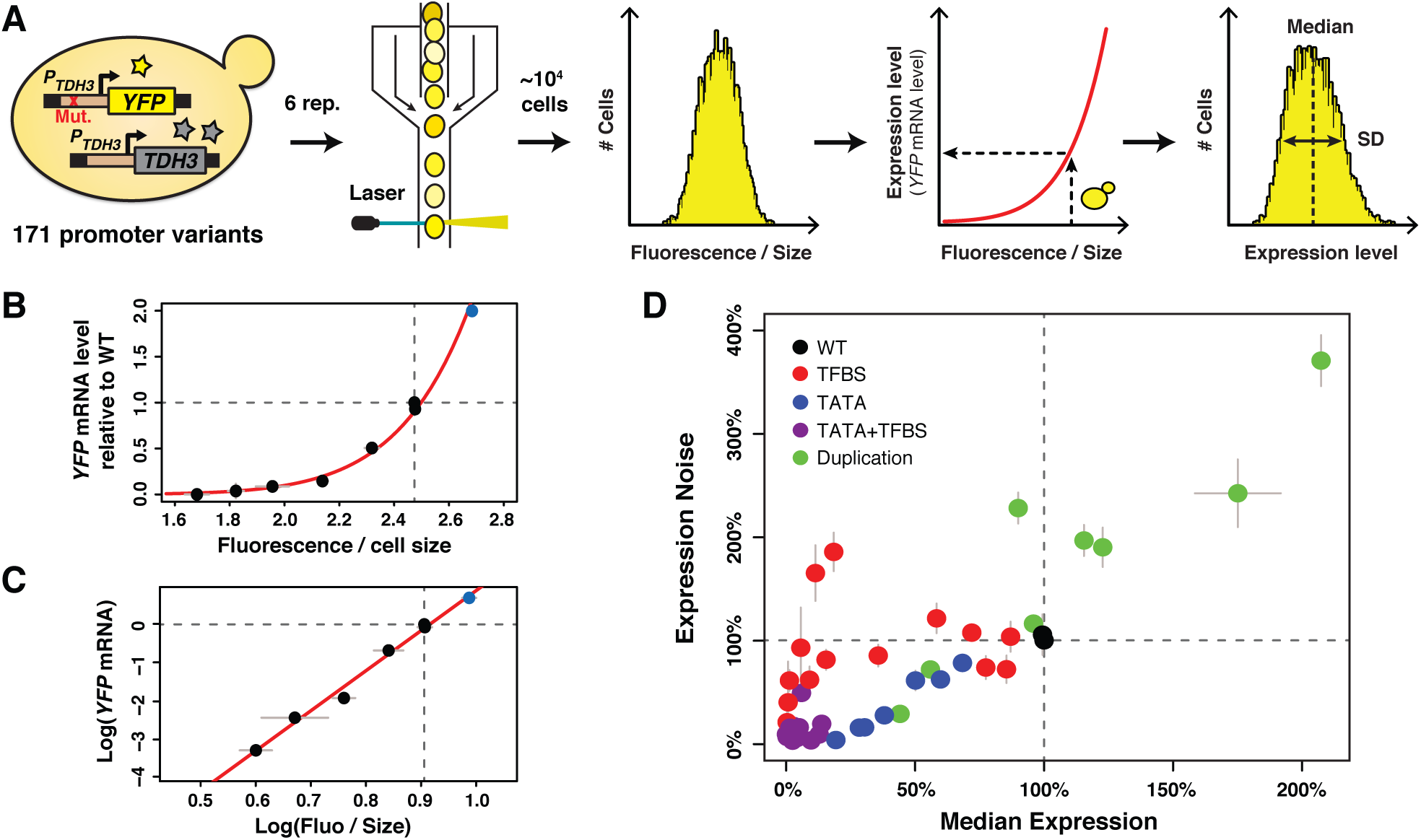
A collection of *TDH3* promoter alleles with incompletely correlated effects on average expression level and expression noise. **(A)** Overview of experimental design used to quantify expression. The transcriptional activity of 171 different variants of the *TDH3* promoter (*P*_*TDH3*_) inserted upstream of the *YFP* coding sequence was quantified using flow cytometry. After growth of six independent samples in rich medium (YPD) for each promoter variant, fluorescence intensity relative to cell size (forward scatter) was measured for ~10,000 individual cells and transformed into *YFP* mRNA estimates using the function shown in **(B)**, allowing characterization of both the median and the standard deviation of expression of the reporter gene. **(B)** Non-linear relationship between *YFP* mRNA level and fluorescence intensity divided by cell size measured on a BD Accuri C6 flow cytometer. **(C)** Linear relationship between the logarithm of *YFP* mRNA level and the logarithm of fluorescence intensity divided by cell size. **(B-C)** *YFP* mRNA level was quantified by pyrosequencing and fluorescence intensity by flow cytometry in three biological replicates of eight strains expressing *YFP* under different variants of *P*_*TDH3*_. Fluorescence intensity was normalized by cell size as described in the Methods section. The red line is the best fit of a function of shape log(*y*) = *a* × log(*x*) + *b* to the data, with *a* = 10.469 and *b* = −9.586. The blue dot represents a strain with two copies of the wild type *P*_*TDH3*_-*YFP* reporter. **(D)** Median expression level and expression noise (noise strength: variance divided by median expression) for 43 *P*_*TDH3*_ alleles. These alleles were chosen to cover a broad range of median expression level and expression noise with an incomplete correlation between these two parameters. Colors represent different types of promoter mutations. **(B-D)** Dotted lines show the activity of the wild type *TDH3* promoter. Error bars are 95% confidence intervals calculated from at least four replicates for each genotype and are only visible when larger than dots representing data.

Effects of 236 point mutations in the *TDH3* promoter, including mutations in RAP1 and GCR1 transcription factor binding sites (TFBS), have previously been described that cause a wide range of average expression levels and expression noise values (Metzger *et al.* 2015). But average expression level and expression noise strongly co-vary among these alleles (Metzger *et al.* 2015), making them insufficient for separating the effects of changes in average expression level and expression noise on fitness. We therefore sought to construct additional promoter alleles that showed a different relationship between average expression level and expression noise. First, we inserted a recognition motif for the GCN4 transcription factor at ten different positions in the *TDH3* promoter because this TFBS was previously found to affect the relationship between expression level and expression noise (Sharon *et al.* 2014). However, the insertion of GCN4 binding sites into *P*_*TDH3*_ did not show the expected departure from the relationship between expression level and expression noise observed for mutations in GCR1 and RAP1 TFBS (Figure 1 - supplementary figure 1). We next mutated the *P*_*TDH3*_ TATA box because previous studies showed that TATA box mutations confer lower expression noise for a given expression level when compared to other types of promoter alterations (Blake *et al.* 2006; Mogno *et al.* 2010; Hornung *et al.* 2012). We generated 112 alleles of the *TDH3* promoter that had between one and five random mutations in the TATA box sequence, which caused the expected lower levels of expression noise than TFBS mutant alleles with similar average expression levels (Figure 1D). We then combined mutations in the TATA box, GCR1 TFBS and/or RAP1 TFBS to further increase the range of expression phenotypes. Finally, we constructed alleles containing two tandem copies of the *P*_*TDH3*_-*YFP* reporter gene with or without mutations in the *P*_*TDH3*_ sequence to sample expression levels closer to and above the wild-type allele.

From a collection of 171 *TDH3* promoter alleles (Table S1, Figure 1 - supplementary figure 1), we selected 43 alleles (Table S2) to study the fitness effects of changes in average expression level and expression noise of the native *TDH3* gene. The average expression level conferred by these 43 *P*_*TDH3*_ alleles (including the wild type allele of *P*_*TDH3*_) ranged from 0% to 207% of the wild type allele and the expression noise ranged from 3% to 371% of the wild type allele (Figure 1D). Most importantly, this set of alleles showed variation in expression noise independent of expression level at expression levels between 0% and 125% of the wild type allele (Figure 1D).

### Fitness effects of changing average TDH3 expression level

To measure the fitness effects of changing *TDH3* expression, we introduced each of these 43 *P*_*TDH3*_ alleles upstream of the *TDH3* coding sequence at the native *TDH3* locus and performed competitive growth assays similar to those described in Duveau *et al.* (2017) (Figure 2A). For each of the eight *P*_*TDH3*_ alleles that contained a duplication of the *P*_*TDH3*_-*YFP* reporter gene, we created a duplication of the entire *TDH3* gene with the two corresponding *P*_*TDH3*_ alleles. We also included a strain with a deletion of the promoter and coding sequence of *TDH3* to sample a *TDH3* expression level of 0%. Prior studies have found that deletion of *TDH3* causes a moderate decrease in fitness in glucose-based media: −5% in Pierce *et al.* (2007) and −6.8% in Duveau *et al.* (2017). Each strain tested was marked with YFP and grown competitively for 30 hours (~21 generations) with a reference genotype marked with a green fluorescent protein (GFP) (Figure 2A). Competitive fitness was determined from the rate of change in genotype frequencies over time and averaged across at least six replicate populations for each genotype tested (Figure 2B). The relative fitness of each strain was then calculated by dividing the competitive fitness of that strain by the competitive fitness of the strain with the wild type allele of *TDH3* (Table S2). This protocol provided a measure of fitness with an average 95% confidence interval of 0.2%. We then related these measures of relative fitness to the expression driven by these *P*_*TDH3*_ alleles of the reporter gene at the *HO* locus, which was found to provide a reliable readout of *P*_*TDH3*_ activity at the native *TDH3* locus for both average expression level and expression noise (Figure 2 – figure supplement 1).

**Figure 2.**
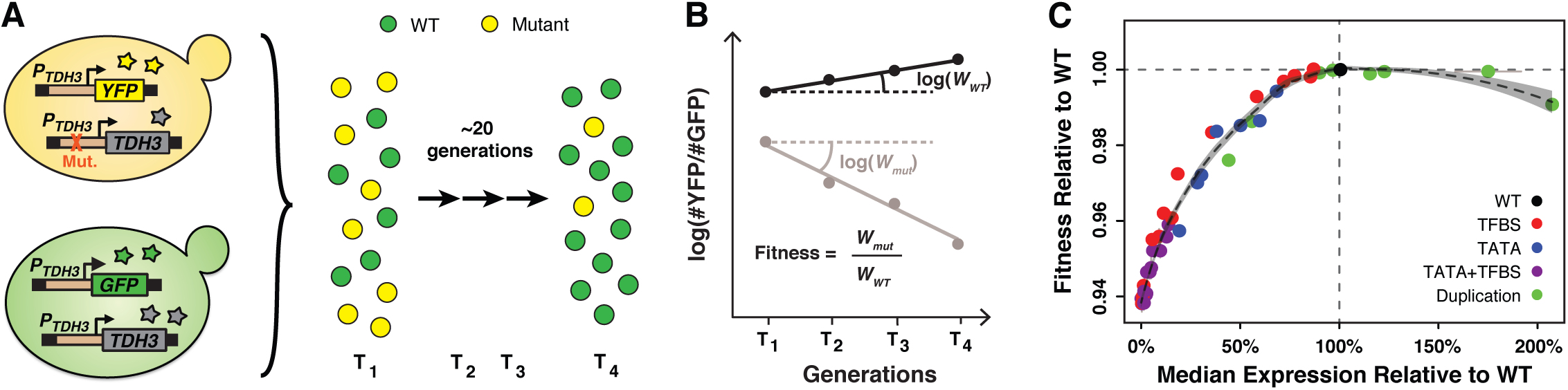
Fitness consequences of variation in *TDH3* expression level. **(A)** Overview of competition assays used to quantify fitness. The 43 *P*_*TDH3*_ alleles whose activity was described in Figure 1D were introduced upstream of the native *TDH3* coding sequence in a genetic background expressing YFP under control of the wild type *TDH3* promoter. A minimum of six replicate populations of the 43 strains were competed for ~20 generations in rich medium (YPD) against a common reference strain expressing GFP under control of the wild type *TDH3* promoter. The relative frequency of cells expressing YFP or GFP was measured every ~7 generations using flow cytometry. **(B)** Competitive fitness was calculated from the change in genotype frequency over time. The relative fitness of each strain was calculated as the mean competitive fitness of that strain across replicates divided by the mean competitive fitness of the strain carrying the wild type allele of *TDH3*. **(C)** Relationship between median expression level of *TDH3* and fitness in rich medium (YPD). Dots show the average median expression and average relative fitness measured among at least four replicates for each of the 43 *P*_*TDH3*_ alleles. Colors represent different types of promoter mutation. Error bars are 95% confidence intervals and are only visible when larger than dots. The dotted curve is the best fit of a LOESS regression of fitness on median expression, using a value of 2/3 for the parameter *α* controlling the degree of smoothing. The shaded area shows the 99% confidence interval of the LOESS fit.

A local regression (LOESS) of average expression level on fitness for the 43 *TDH3* alleles and the *TDH3* deletion showed a non-linear relationship with a plateau of maximal fitness near the wild type expression level (Figure 2C) similar to that described in Duveau *et al.* (2017). Deletion of *TDH3* (expression level of 0% in Figure 2C) caused a statistically significant decrease in fitness of 6.1% relative to the wild type allele (t-test, P = 6.4 x 10^−10^). The minimum change in *TDH3* expression level that significantly impacted fitness was a 14.6% decrease in average expression relative to wild type, which reduced fitness by 0.19% (*t*-test, *P* = 0.00045). Overexpressing *TDH3* up to 175% did not significantly impact growth rate, but the 207% expression level of the strain carrying a duplication of the wild type *TDH3* gene caused a 0.92% reduction in fitness (Figure 2C; *t*-test, *P* = 1.4 x 10^−7^). Notably, none of the 42 mutant alleles of *TDH3* conferred a significantly higher fitness than the wild type allele (one-sided *t*-tests, P > 0.05), indicating that the wild type expression level of *TDH3* is near an optimum for growth in the environment assayed.

### Disentangling the effects of TDH3 expression level and expression noise on fitness

Residual variation was observed around the LOESS fitted line relating expression level to fitness (Figure 2C) that we hypothesized might be explained by differences in noise among genotypes. To examine the effects of differences in expression noise on fitness independent of differences in average expression level, we used the residuals from a local regression of expression noise on expression level for the alleles with average expression levels between 0% and 125% of the wild type allele to define a metric called ΔNoise (Figure 3A; Figure 3 – figure supplement 1A-D). This metric was not significantly correlated with expression level (Figure 3 – figure supplement 2). *TDH3* alleles with positive ΔNoise values had a higher level of noise than expected based on their expression level and were classified as “high noise”, whereas *TDH3* alleles with negative ΔNoise values had lower levels of noise than expected given their expression level and were classified as “low noise”.

**Figure 3.**
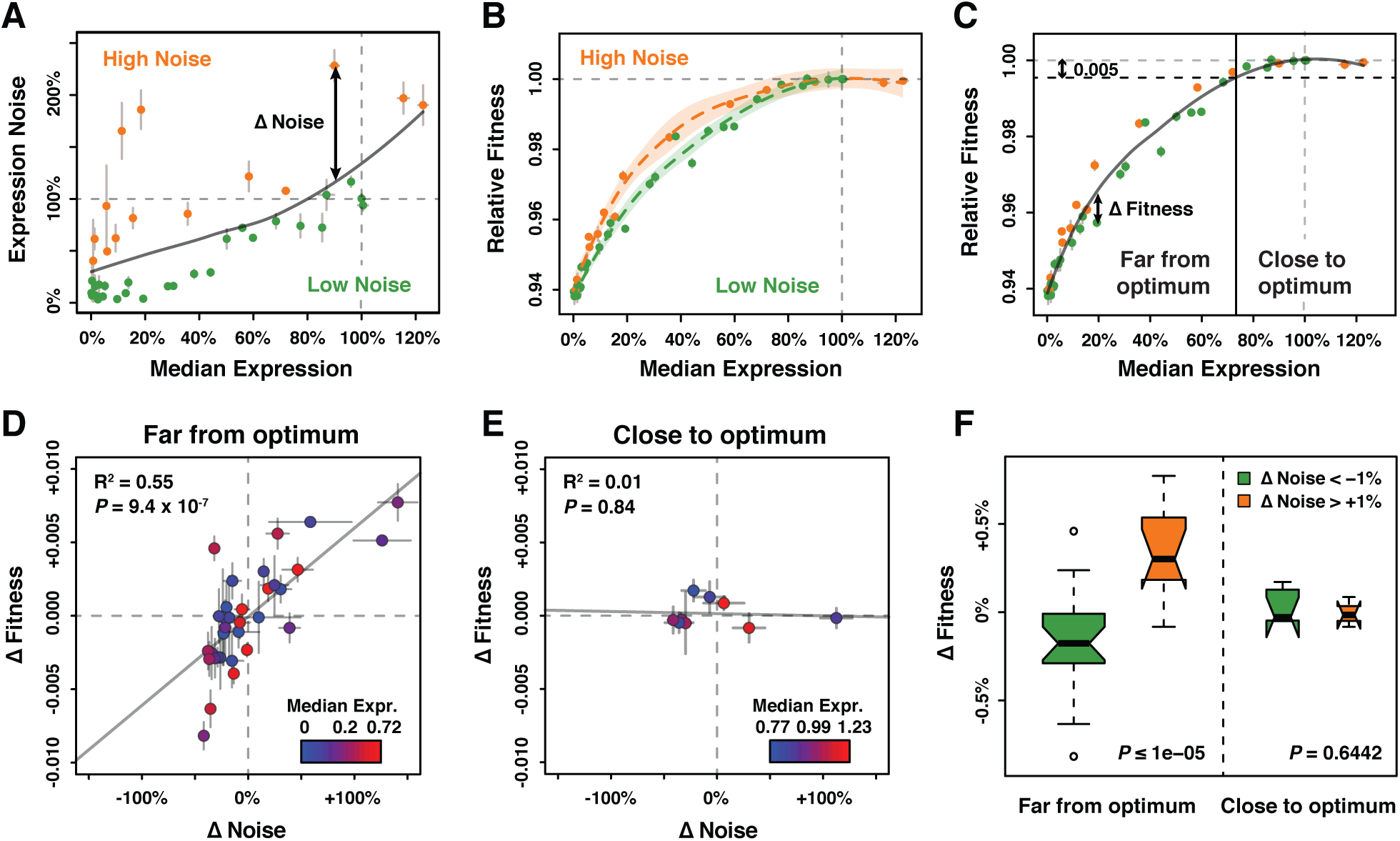
Effect of *TDH3* expression noise on fitness. **(A)** Separation of the 43 *P*_*TDH3*_ alleles into two categories based on their effects on median expression level and expression noise (noise strength). The gray curve shows the LOESS regression of noise on median expression using a value of 2/3 for the smoothing parameter. Data points falling below the curve (green) correspond to *P*_*TDH3*_ alleles with low noise given their median level of activity. Data points above the curve (orange) correspond to *P*_*TDH3*_ alleles with high noise given their median activity. The residual of the LOESS regression (“Δ Noise”) is a measure of noise independent of median expression. **(B)** Relationships between median expression level and fitness for strains with low noise (green, Δ Noise < −1%) and high noise (orange, Δ Noise > +1%). The two LOESS regressions were performed with smoothing parameter *α* equal to 2/3. **(C)** Partition of *P*_*TDH3*_ alleles into two groups based on the distance of their median activity to the optimal level of *TDH3* expression. The expression optimum (vertical gray dotted line) corresponds to the expression level predicted to maximize fitness from the LOESS regression of fitness on median expression (gray curve). The expression level at which the predicted fitness is 0.005 below the maximal fitness was chosen as the threshold (vertical black dotted line) separating promoters with median activity “close to optimum” from promoters with median activity “far from optimum”. The residual of the LOESS regression (“Δ Fitness”) is a measure of fitness independent of the median *TDH3* expression level. Dots are colored as in **(B)**. **(D)** Relationship between Δ Noise and Δ Fitness when median expression is far from optimum. **(E)** Relationship between Δ Noise and Δ Fitness when median expression is close to optimum. **(D-E)** The best linear fit between Δ Noise and Δ Fitness is shown as a gray line, with the coefficient of determination (“R^2^”) and the significance of the Pearson’s correlation coefficient (“*P*”) indicated in the upper left of each panel. Dots are colored based on median expression levels of the corresponding *P*_*TDH3*_ alleles as indicated by color gradient. **(A-E)** Error bars show 95% confidence intervals calculated from at least four replicate samples and are only visible when larger than symbols representing data points. **(F)** Comparison of Δ Fitness between genotypes with low noise strength (green, Δ Noise < −1%) and genotypes with high noise strength (red, Δ Noise > +1%). Thick horizontal lines represent the median Δ Fitness among genotypes and notches display the 95% confidence interval of the median. Bottom and top lines of each box represent 25^th^ and 75^th^ percentiles. Width of boxes is proportional to the square root of the number of genotypes included in each box. Permutation tests were used to assess the significance of the difference in median Δ Fitness between genotypes with low and high noise and *P*-values are shown in lower right corners.

We then compared the relationship between expression level and fitness for genotypes in the high noise and low noise classes (Figure 3B). We found that promoter alleles with positive ΔNoise tended to show a higher fitness than strains with negative ΔNoise (Figure 3B, Figure 3 – figure supplement 1E-H). This beneficial effect of noise on fitness was surprising given prior evidence that selection favored alleles of *P*_*TDH3*_ with low expression noise in natural populations (Metzger *et al.* 2015). We noticed, however, that the fitness benefit of increasing expression noise was limited to a particular range of average expression levels. Specifically, positive ΔNoise was associated with higher fitness only for average expression levels between 2% and 80% of the wild type expression level (Figure 3B). Above 80% of expression, no clear difference in fitness was detected between strains with positive and negative ΔNoise (Figure 3B). These same trends were also observed for the other metrics of noise (Figure 3 – figure supplement 1E-H).

Based on these observations and prior theoretical work (Tǎnase-Nicola and Ten Wolde 2008), we hypothesized that the distance between the average expression level of a *P*_*TDH3*_ allele and the optimal level of *TDH3* expression influenced how a change in expression noise impacted fitness. To test this hypothesis, the 43 promoter alleles were split into two groups depending on the distance of their average expression level from the optimal expression level of *TDH3*. Using a local regression of fitness on average expression level, we inferred the value of average expression that would confer a fitness reduction of 0.5% from maximal fitness. Promoter alleles for which the median activity was below this threshold were considered to be “far from optimum” and promoter alleles with median activity above the threshold were considered to be “close to optimum” (Figure 3C). A metric called ΔFitness, corresponding to the residuals of a local regression of fitness on average expression, was calculated to remove the confounding effect of average expression levels on fitness (Figure 3C, Figure 3 – figure supplement 1I-L, Figure 3 – figure supplement 3). We found that changes in noise (ΔNoise) and changes in fitness (ΔFitness) were positively correlated for genotypes classified as far from the optimum (Pearson correlation coefficient: r = 0.74, *P* = 9.36 x 10^−7^, Figure 3D, figure supplement 4A-D), but not for genotypes classified as close to the optimum (r = −0.08, *P*= 0.84, Figure 3E, figure supplement 4E-H). This result was robust to variation in the choice of the smoothing parameter used for the local regression of noise on average expression, the choice of the smoothing parameter used for the local regression of fitness on average expression, and the fitness threshold used to separate strains with expression levels close and far from optimum (Figure 3 – figure supplement 5, 6, 7).

As an alternative way to test for the impact of expression noise on fitness, we compared ΔFitness for genotypes with positive and negative values of ΔNoise. Permutation tests were used to assess the significance of differences in ΔFitness by randomly shuffling values of ΔNoise among genotypes. Consistent with the correlation analyses, genotypes with positive ΔNoise showed a significantly greater median value of ΔFitness than genotypes with negative ΔNoise at expression levels far from optimum (10^5^ permutations, *P*_*ΔNoise*_ ≤ 10^−5^; Figure 3F). Among genotypes with average expression close to optimum, no significant difference in median ΔFitness was detected between the positive and negative ΔNoise groups (10^5^ permutations, *P*_*ΔNoise*_ = 0.6442) (Figure 3F). The same pattern was observed for all metrics of noise and was not driven by differences in average expression levels between the two ΔNoise groups (Figure 3 – figure supplement 8).

### Direct measurements of the effect of expression noise on relative fitness

The results presented in the preceding section provide strong evidence that variation in *TDH3* expression can directly affect fitness, but the methods used have at least two limitations. First, ΔNoise and ΔFitness values can be influenced by the set of *P*_*TDH3*_ alleles included in the analyses since they are regression residuals. Second, comparisons of fitness among *P*_*TDH3*_ genotypes rely upon the assumption that fitness effects are transitive, *i.e.* that differences in fitness between two strains are accurately reported by competitive growth against a third reference strain. Even though such transitivity has often been verified (De Visser and Lenski 2002; Elena and Lenski 2003; Gallet *et al.* 2012), intransitive competition has been observed in several organisms, including yeast (Paquin and Adams 1983). To test whether differences in *TDH3* expression noise affect fitness without calculating regression residuals and without assuming transitivity, we performed direct competition assays between strains with *P*_*TDH3*_ promoter alleles that showed similar average expression levels but different levels of expression noise.

Five pairs of *TDH3* alleles for which (i) the median level of activity was similar between the two promoters of each pair, (ii) the level of noise was different between the two promoters of each pair, and (iii) the median level of activity varied among different pairs were chosen from the full set of 171 alleles described above (Figure 4A; Table S3). The promoter variants of four of these pairs were included in the indirect competition assays and showed the general pattern of increased fitness with increased expression noise when average expression was far from optimum and no significant difference in fitness despite differences in expression noise when average expression was close to optimum (Figure 4B). Promoters in the fifth pair were not among the 43 alleles included in the indirect competition experiment but were selected for the direct competition assays because they showed variation in expression noise at average expression levels close to wild type (purple points in Figure 4A).

**Figure 4.**
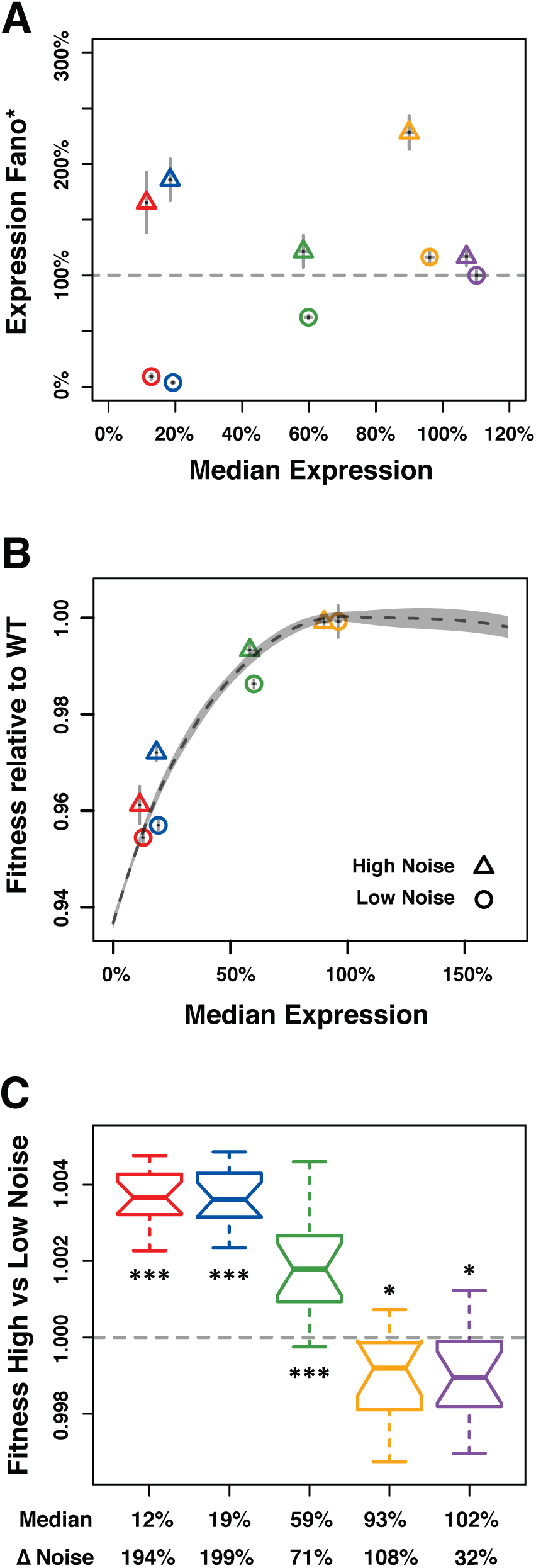
Direct competition between genotypes with different levels of noise but similar median levels of *TDH3* expression in glucose. **(A-C)** Different colors are used to distinguish pairs of genotypes (*P*_*TDH3*_ alleles) with different median expression levels. **(A)** Median expression level and expression noise (noise strength) for five pairs of genotypes that were competed against each other. Each pair comprises one genotype with low expression noise (circle) and one genotype with high expression noise (triangle). **(B)** Relative fitness for four pairs of genotypes measured in competition assays against the common GFP reference strain. One pair is missing (purple in **(A)**) because the corresponding *P*_*TDH3*_ alleles were not part of the 43 alleles included in the initial competition experiment. **(A-B)** Error bars show 95% confidence intervals obtained from at least three replicates. **(C)** Competitive fitness of high noise strains relative to low noise strains measured from direct competition assays. Each box represents fitness data from 16 replicate samples. Thick horizontal lines represent the median fitness across replicates and notches display the 95% confidence interval of the median. The bottom and top lines of each box represent 25^th^ and 75^th^ percentiles. Statistical difference from a fitness of 1 (same fitness between the two genotypes) was determined using *t*-tests (*: 0.01 < *P* < 0.05; **: 0.001 < *P* < 0.01; ***: *P* < 0.001).

For each of the five pairs, the low noise genotype and the high noise genotype were directly competed against each other under the same conditions used in the competitive growth fitness assay described above except that we doubled the number of generations and the number of replicates to increase the sensitivity of our fitness estimates. In addition, we used pyrosequencing (Neve *et al.* 2002) instead of flow cytometry to determine the relative frequency of the two genotypes at each time point because the two strains competed against each other could not be distinguished based on fluorescence. Relative fitness of the high and low noise genotypes in each pair was calculated based on the changes in relative allele frequency during competitive growth.

For the three pairs of genotypes with an average expression level far from optimum (12%, 19%, and 59% average expression relative to wild type), fitness of the high noise genotype relative to the low noise genotype was significantly greater than 1 (Figure 4C), indicating that the high noise genotype grew faster than the low noise genotype. This result was consistent with the differences in fitness measured from the indirect competition assays (Figure 4B). By contrast, both pairs of strains with an average expression level closer to the fitness optimum (93% and 102% relative to wild type expression levels) showed slightly but significantly lower fitness of the high noise genotype than the low noise genotype (Figure 4C). In these cases, higher expression noise resulted in a ~0.1% decrease in fitness relative to lower noise. This difference was detectable with the direct competition assay because the average span of the 95% confidence intervals of fitness estimates was 0.1%, which is half of the 0.2% average 95% confidence intervals from the indirect competition assay described above.

Taken together, our empirical measures of relative fitness show that higher expression noise for *TDH3* is beneficial when average expression level is far from the optimum, but deleterious when average expression is close to the optimum. An intuitive explanation of this phenomenon is that when the average expression level is close to the optimum, increasing expression noise can result in enough cells with suboptimal expression to decrease fitness of the population. Conversely, when the average expression level is far from the optimum, increasing expression noise can result in enough cells with expression closer to the optimum to increase fitness of the population. These effects of expression noise on population fitness can result from differences in expression level among cells causing differences in the cell division rate (a.k.a. doubling time) among cells (Kiviet *et al.* 2014). To better understand the interplay among average expression level, expression noise, and fitness, we developed a simple computational model that allowed us to (1) vary the expression mean and noise independently while holding all other parameters constant, (2) track the resulting single cell growth dynamics, and (3) evaluate the consequences for population fitness.

### Simulating population growth reveals fitness effects of noise

To further investigate how the distribution of expression levels among genetically identical cells influences population fitness, we modeled the growth of clonal cell populations that differed in the mean expression level and expression noise for a single gene. In this model, we specified a function defining the relationship between the expression level of a cell and the doubling time of that cell. Following each cell division, the expression levels of mother and daughter cells were sampled independently from an expression distribution characterized by its mean and noise (Figure 5A). The doubling time of each cell was then calculated from its expression level (Figure 5B). Each clonal population was allowed to expand for the same amount of time, increasing in size at a rate determined by the doubling times of the cells sampled (Figure 5C). Competitive fitness was then determined from the population size obtained at the end of each simulation experiment relative to the population size obtain for a constant “wild type” competitor (Figure 5D, Table S4). 100 independent simulations were performed for each unique combination of mean expression level and expression noise. Three metrics of expression noise were used for this work: noise strength (similar to Fano factor, Figure 6), standard deviation (Figure 6 – figure supplement 1A,C) and coefficient of variation (Figure 6 – figure supplement 1B,D).

**Figure 5.**
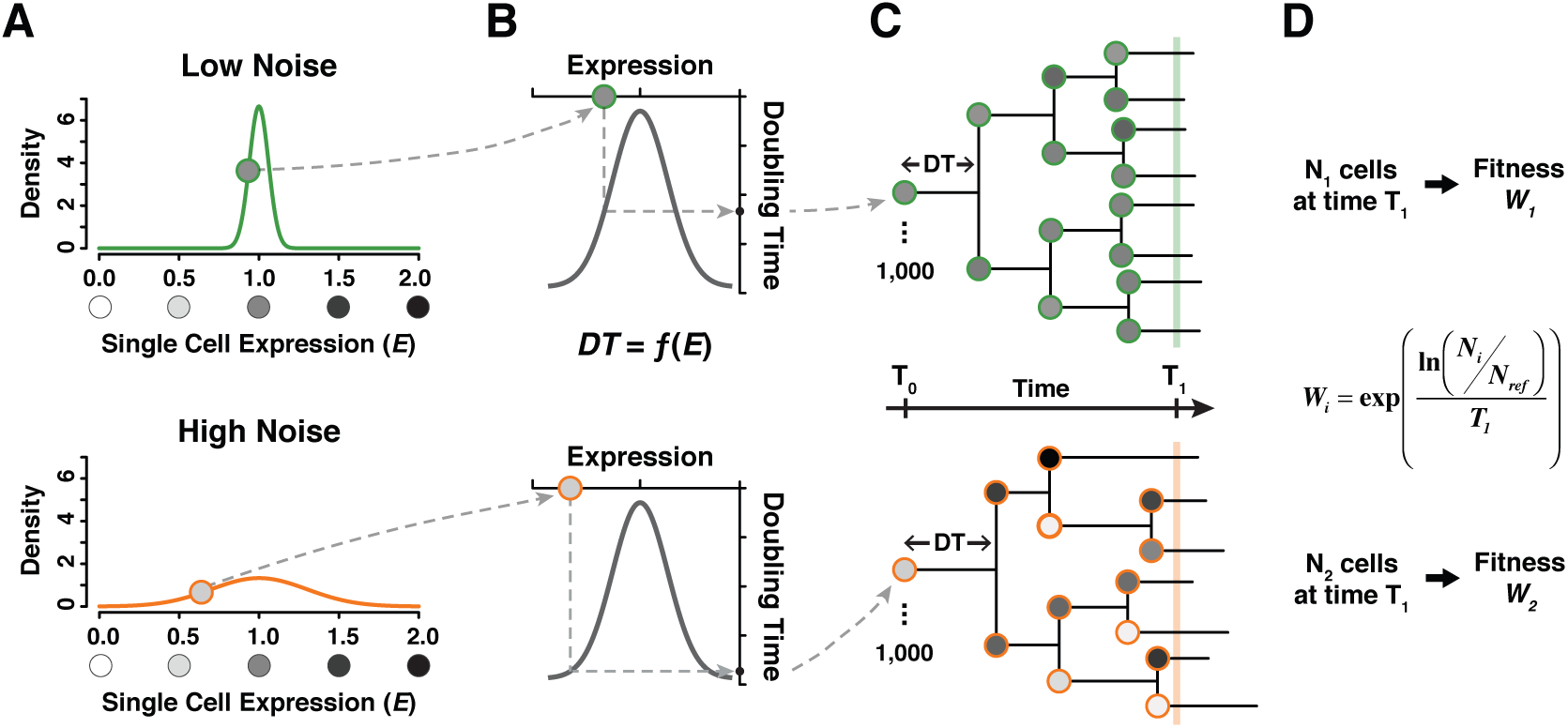
A simple model linking single cell expression levels to population fitness. **(A)** In our model, the expression level *E* of individual cells is randomly drawn from a normal distribution 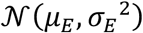 is lower for a genotype with low expression noise (top, green line) and higher for a genotype with high expression noise (bottom, orange line). **(B)** The doubling time *DT* of individual cells is directly determined from their expression level using a function *DT* = *f*(*E*). **(C)** The growth of a cell population is simulated by drawing new values of expression converted into doubling time after each cell division. In this example, doubling time is more variable among cells for the population showing the highest level of expression noise. **(D)** Population growth is stopped after a certain amount of time (1000 minutes in our simulations) and competitive fitness is calculated from the total number of cells produced by the tested genotype relative to the number of cells in a reference genotype with *μ*_*E*_ = 1 and *θ*_*E*_ = 0.1. In this example, fitness is lower for the genotype with higher expression noise (bottom) because it produced less cells than the genotype with lower expression noise (top).

**Figure 6.**
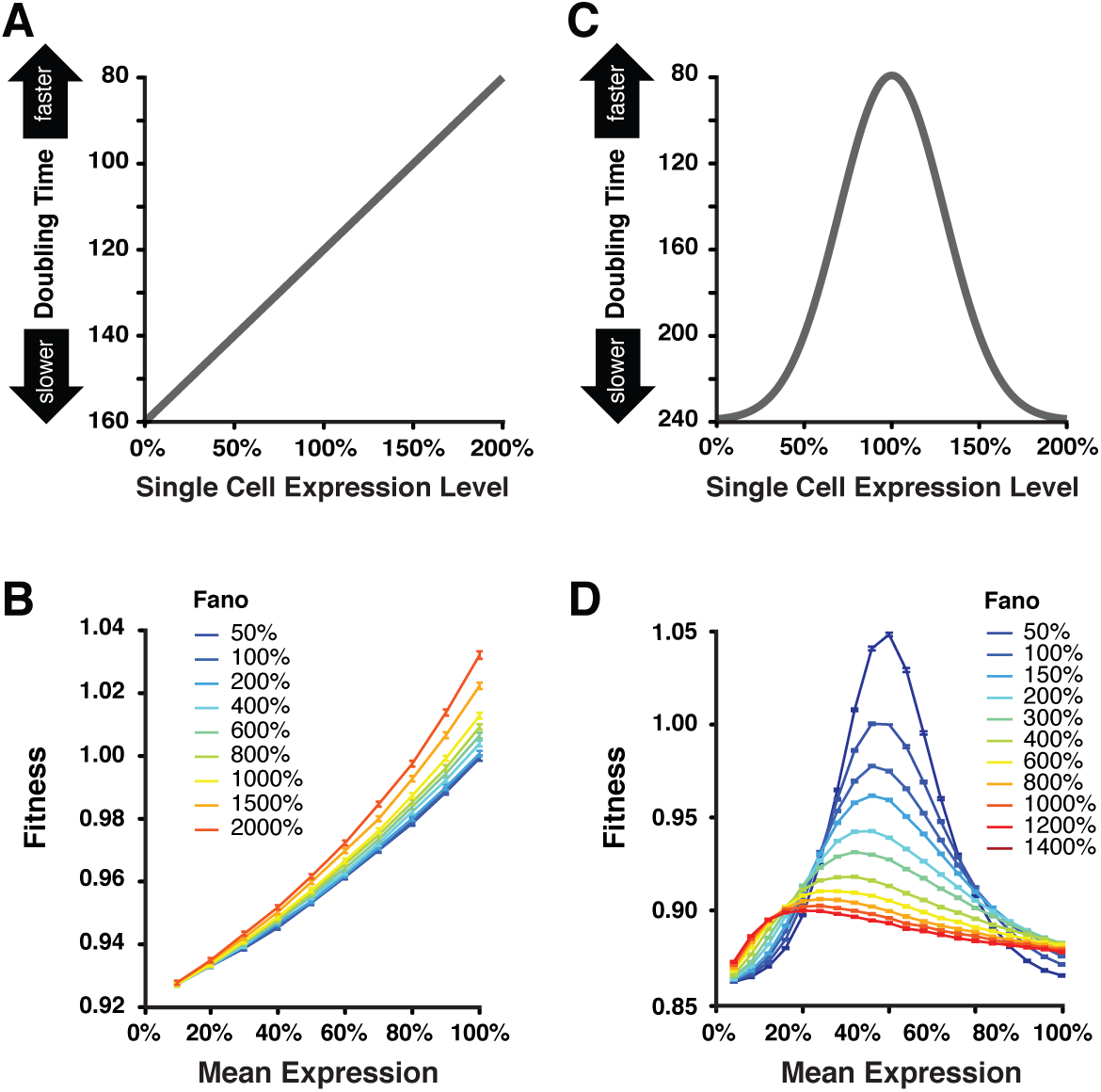
Simulating the effect of expression noise on fitness at different median expression levels. **(A)** The linear function *DT* = −40 × *E* + 160 relating the expression level of single cells to their doubling time used for the first set of simulations. **(B)** Relationship between mean expression (*μ*_*E*_) and fitness at nine values of expression noise (noise strength: *θ*_*E*_^2^/*μ*_*E*_) ranging from 50% to 2000% using the linear function shown in **(A)**. **(C)** Gaussian function TW = −160 × exp(−(*E* − 1)^2^/0.18) + 240 relating the expression level of single cells to their doubling time used in the second set of simulations. This function shows an optimal expression level at *E* = 1, where doubling time is minimal (*i.e.*, fastest growth rate). **(D)** Relationship between mean expression (*μ*_*E*_) and fitness at 11 values of expression noise (noise strength: *θ*
_*E*_^2^/*μ*_*E*_) ranging from 50% to 1400% using the Gaussian function shown in **(C)**. **(B,D)** Error bars show 95% confidence intervals of mean fitness calculated from 100 replicate simulations for each combination of mean expression and expression noise values.

To calculate doubling times from single cell expression levels, we first used a linear function akin to directional selection in which increases in expression level resulted in shorter doubling times (faster growth) (Figure 6A). With this relationship, higher levels of expression noise conferred higher population fitness for a given mean expression level (Figure 6B), a pattern more pronounced for high values of mean expression and observed for all metrics of noise (Figure 6 – figure supplement 1A,B). This finding is consistent with prior work demonstrating that an increased variability of doubling time among individual cells is sufficient to increase fitness at the population level (Tǎnase-Nicola and Ten Wolde 2008; Cerulus *et al.* 2016; Hashimoto *et al.* 2016; Nozoe *et al.* 2017). This is because the doubling time of a population tends to be dominated by the doubling time of the fastest dividing cells in the population, *i.e.* population doubling time is higher than the mean doubling time among all cells in the population.

Next, we used a Gaussian function akin to stabilizing selection in which an intermediate expression level produced the shortest doubling time (faster growth), while lower or higher expression than this optimum would increase doubling time (slower growth) (Figure 6C). With this function, we found that the fitness effects of increasing expression noise depended on the mean expression level. Specifically, increasing expression noise increased fitness when the average expression level was far from the optimal expression level and it decreased fitness when the average expression level was close to the optimum (Figure 6D), similar to the pattern we observed with our empirical fitness data and in agreement with theoretical work by Tǎnase-Nicola and Ten Wolde (2008). This result was observed for all three metrics of noise, suggesting it is robust to the different scaling relationships between the mean expression level and variability around the mean captured by different metrics of noise (Figure 6 – figure supplement 1C,D).

These *in silico* analyses not only provide a plausible mechanistic explanation for our empirical finding that increasing noise can be both beneficial and deleterious in a single environment but they also show that increasing expression noise can alter the effects of changes in mean expression level on fitness. Specifically, when expression noise is high (red lines on Figure 6D and Figure 6 – figure supplement 1C,D), changes in mean expression level are predicted to have much smaller impacts on fitness than equivalent changes when expression noise is low (blue lines on Figure 6 and Figure 6 – figure supplement 1C,D). This pattern is also readily apparent when changes in expression noise, instead of changes in mean expression level, are plotted as a function of population fitness (Figure 6 - figure supplement 2). These observations are consistent with a previously published population genetic model showing that increasing expression noise can reduce the efficacy of natural selection acting on mean expression level (Wang and Zhang 2011).

## Conclusions

Despite many studies providing evidence that natural selection can (Tǎnase-Nicola and Ten Wolde 2008; Wang and Zhang 2011; Barroso *et al.* 2018) and has (Fraser *et al.*2004; Lehner 2008; Zhang *et al.* 2009; Metzger *et al.* 2015) acted on expression noise, the precise effects of expression noise on fitness have proven difficult to measure empirically. This difficulty arises from the facts that (1) most mutations that alter expression noise also alter mean expression in a correlated fashion, making it difficult to isolate the effects of changes in expression noise on fitness (Hornung *et al.* 2012; Keren *et al.* 2016; Liu *et al.* 2016), and (2) the magnitude of fitness effects resulting from changes in expression noise is expected to be smaller than that resulting from changes in mean expression level (Zhang *et al.* 2009). In this study, we overcame these challenges by surveying a broad range of promoter alleles for their effects on mean expression level and expression noise, measuring the fitness effects of a subset of these alleles with reduced dependency between effects on mean expression level and expression noise, and using an assay for fitness with power to detect changes as small as 0.1%. We found that the fitness effects of changes in expression noise are indeed generally much smaller than changes in expression level, although they are large enough to be acted on by natural selection in wild populations of *S. cerevisiae* (Wagner 2005; Metzger *et al.* 2015).

We also show that changes in expression noise can be beneficial or deleterious depending on the distance between the mean expression level and the expression level conferring optimal fitness in the environment examined, with increases in expression noise deleterious near the optimal expression level, consistent with data for *TDH3* in Metzger *et al.* 2015. Although our empirical work focused solely on the *TDH3* gene, the small number of parameters in our simulation model producing the same pattern as these empirical data suggests that the observed relationship among fitness, average expression level and expression noise are likely generalizable to other genes. That said, the precise the relationship between expression noise and fitness at the population level is expected to be shaped by the relationship between average expression level and doubling time of single cells as well as the temporal dynamics of expression in single cells (Blake *et al.*2006; Tǎnase-Nicola and Ten Wolde 2008). These single-cell properties will thus be important to characterize in future work using tools such as time-lapse microscopy.

Assuming that the average expression level of a population is near the fitness optimum in a stable environment, but further from the optimum following a change in the environment, our results unify studies showing that increasing expression noise tends to be deleterious in a constant environment but beneficial in a fluctuating one (Fraser *et al.* 2004; Blake *et al.* 2006; Batada and Hurst 2007; Lehner 2008; Tǎnase-Nicola and Ten Wolde 2008; Zhang *et al.* 2009; Fraser and Kærn 2009; Ito *et al.* 2009; Wang and Zhang 2011; Levy *et al.* 2012; Wolf *et al.* 2015; Liu *et al.* 2015; Keren *et al.* 2016). Expression noise may be particularly important in the early phase of adaptation to a fluctuating environment, when a new expression optimum makes an increase in noise beneficial and before expression plasticity evolves as a more optimal strategy (Wolf *et al.* 2015; Duveau, Yuan, *et al.* 2017). Perhaps most interestingly, these data show that high levels of noise can transition from being deleterious to beneficial even in a stable environment following a change in average expression level to a level further from the fitness optimum. We note, however, that the often correlated effects of promoter mutations on mean expression level and expression noise (Hornung *et al.* 2012; Carey *et al.* 2013; Sharon *et al.* 2014; Vallania *et al.* 2014) may limit the ability of natural selection to optimize both mean expression level and expression noise. Future work investigating the effects of other types of mutations on mean expression level, expression noise, and fitness in multiple environments is needed to define more fully the range of variation affecting gene expression upon which natural selection can act.

## Materials and Methods

### Yeast strains: genetic backgrounds

All strains used in this work were haploids with similar genetic backgrounds that were derived from crosses between BY4724, BY4722, BY4730 and BY4742 (Brachmann *et al.* 1998) and carry the alleles *RME1(ins-308A); TAO3(1493Q)* from Deutschbauer and Davis (2005) and *SAL1; CAT5(91M); MIP1(661T)* from Dimitrov *et al.* (2009) that contribute to increased sporulation efficiency and decreased petite frequency relative to the alleles of the laboratory S288c strain. The construction of this genetic background is described in more detail in Metzger *et al.* (2016). Strains used to assay transcriptional activity and fitness (described in detail below) had different mating types and drug resistance markers, but these differences did not significantly affect *P*_*TDH3*_ transcriptional activity (Figure 2 – figure supplement 2A,B).

### Yeast strains: construction of strains used to assay transcriptional activity

Transcriptional activity (average expression level and expression noise) was assayed for 171 *P*_*TDH3*_ alleles in *S. cerevisiae* strains carrying a fluorescent reporter construct inserted at the *HO* locus on chromosome IV in *MATα* cells (Metzger et al, 2016). From these alleles (Table S1), 43 were selected for assaying fitness effects of changing *TDH3* expression (Table S2). 36 of the final 43 *P*_*TDH3*_ alleles carried a single copy of a reporter construct consisting of the *TDH3* promoter followed by the *Venus YFP* coding sequence, the *CYC1* terminator and an independently transcribed *KanMX4* drug resistance cassette Metzger *et al.* (2016). 7 of the final 43 *P*_*TDH3*_ alleles variants consist of two copies of the *P*_*TDH3*_-*YFP*-*T*_*CYC1*_ construct in tandem separated by a *URA3* cassette. The different *P*_*TDH3*_ alleles contain mutations located either in the known binding sites for *GCR1* and *RAP1* transcription factors, in the TATA box or in combinations of both, as described below. The wild type allele of *P*_*TDH3*_ consists of the 678 bp sequence located upstream of the *TDH3* start codon in the genome of the laboratory strain S288c, with a single nucleotide substitution that occurred during the construction of the *P*_*TDH3*_-*YFP*-*T*_*CYC1*_ construct (A -> G located 293 bp upstream of the start codon). This substitution is present in all *P*_*TDH3*_ alleles used in this study. The effect of this mutation on *P*_*TDH3*_ activity in YPD medium was previously described (Metzger *et al.* 2015).

#### Single TFBS mutants

A set of 236 point mutations corresponding to almost all C -> T and G -> A substitutions in the *TDH3* promoter was previously inserted upstream of a *YFP* reporter gene on chromosome I in the BY4724 genetic background (Metzger *et al.* 2015). From these, we selected seven *P*_*TDH3*_ alleles for which the transcriptional activity spanned a broad range of median fluorescence levels when cells were grown in glucose medium (25% to 90% relative to wild type expression level). These seven promoters carried mutations either in the GCR1 or RAP1 transcription factor binding sites (TFBS) previously characterized in the *TDH3* promoter (Yagi *et al.* 1994). Each *P*_*TDH3*_ allele was inserted upstream of *YFP* at the *HO* locus using the *dellitto perfetto* approach (Stuckey *et al.* 2011). Briefly, in the reference strain YPW1002 carrying the wild-type *P_TDH3_-YFP-T_CYC1_* construct at *HO* (Metzger *et al.* 2016), we replaced *P*_*TDH3*_ with a *CORE-UH* cassette (*COunterselectable REporter URA3-HphMX4* amplified from plasmid *pCORE-UH* using oligonucleotides 1951 and 1926 in Table S5) to create strain YPW1784. Then, each of the seven *P*_*TDH3*_ alleles was amplified by PCR using oligonucleotides 2276 and 2277 (Table S5) and transformed into YPW1784 to replace the *CORE-UH* cassette and allow expression of *YFP* (Metzger *et al.* 2015). Correct insertion of *P*_*TDH3*_ alleles was verified by Sanger sequencing of PCR amplicons obtained with primers 2425 and 1208 (Table S5).

#### Double TFBS mutants

To sample average expression levels less than 25% of wild type, we created and measured the activity of 12 *P*_*TDH3*_ alleles containing mutations in two different TFBS. We then selected seven of these alleles to be included in the final set of 43 *P*_*TDH3*_ alleles (Table S2). Point mutations from different alleles were combined on the same DNA fragment using PCR SOEing (Splicing by Overlap Extension). First, left fragments of *P*_*TDH3*_ were amplified from genomic DNA of strains carrying the most upstream TFBS mutations. These PCRs used a common forward primer (2425 in Table S5) and a reverse primer containing the most downstream TFBS mutation to be inserted (P4E8, P4E5, P4G8 or P4G7 in Table S5). In parallel, right fragments of *P*_*TDH3*_ were amplified from YPW1002 gDNA using forward primers containing the most downstream TFBS mutations (P1E8, P1E5, P1G8 or P1G7 in Table S5) and a common reverse primer (104 in Table S5). Then, equimolar amounts of the overlapping upstream and downstream fragments of *P*_*TDH3*_ were mixed and 25 PCR cycles were performed to fuse both fragments together and to reconstitute the full promoter. Finally, the fused fragments were further amplified for 35 cycles using oligonucleotides 2425 and 1305 (Table S5) and the final products were transformed in strain YPW1784. The presence of desired mutations in *P*_*TDH3*_ was confirmed by Sanger sequencing of amplicons obtained with primers 1891 and 1208 (Table S5).

#### GCN4 binding sites

To try to create variation in noise independent of the median expression level, we inserted GCN4 binding sites at several locations in the *TDH3* promoter because GCN4 binding sites in synthetic promoters were shown to increase expression noise (CV^2^) relative to average expression level (Sharon *et al.* 2014). We introduced substitutions in *P*_*TDH3*_ to create the GCN4 binding motif TGACTCA at 10 different locations (−121, −152, −184, −253, −270, −284, −323, −371, −407 and −495 upstream of start codon) that originally differed by one, two or three nucleotides from this motif. Targeted mutagenesis was performed using the same PCR SOEing approach as described in Metzger *et al.* (2015) (see Table S5 for the list of primers used to insert GCN4 binding sites) and the resulting PCR products were transformed into strain YPW1784. Correct insertion of the TGACTCA motif was confirmed by Sanger sequencing. However, none of the 10 alleles of *P*_*TDH3*_ with GCN4 binding sites showed the expected increase in expression noise when cells were grown in glucose (Figure 1 - figure supplement 1). This could be due to the genomic context being different from the synthetic library used in Sharon *et al.* (2014) or to the fact that *P*_*TDH3*_ is one of the most highly active promoters in the yeast genome. None of these 10 alleles were included in the set of 43 *P*_*TDH3*_ alleles used for fitness assays.

#### TATA box mutants

A second strategy we employed to create variation in expression noise independent of median expression was to mutate the TATA box in the *TDH3* promoter because the presence of a canonical TATA box in yeast promoters has been associated with elevated expression noise (Newman *et al.* 2006). Mutations in the TATA box were also shown to have a clearly distinct impact on expression noise compared to other types of *cis*- regulatory mutations (Blake *et al.* 2006; Hornung *et al.* 2012). We used a random mutagenesis approach to create a large number of alleles with one or several mutations in the *P*_*TDH3*_ TATA box. Variants were obtained using PCR SOEing as described above, except that one of the internal overlapping oligonucleotides (primer 2478, Table S5) used to amplify the downstream fragment of *P*_*TDH3*_ contained a degenerate version of the wild type TATA box (TATATAAA at position −141 upstream of start codon). This oligonucleotide was synthesized by Integrated DNA Technologies using hand-mixed nucleotides at the eight bases of the TATA box with a proportion of 73% of the wild type nucleotide and 9% of each of the 3 alternative nucleotides. At this level of degeneracy, ~10% of the DNA molecules should carry no mutation, ~25% should carry a single mutation in the TATA box, ~35% two mutations, ~20% three mutations and ~10% four mutations or more. The degenerate primer was used with oligonucleotide 104 to amplify the right fragment of *P*_*TDH3*_, and the overlapping primer 2479 was used with oligo 2425 to amplify the left fragment (Table S5). Then, these fragments were fused and amplified as described above for the TFBS mutants. Six independent transformations of the fragments containing random mutations in the TATA box were performed in strain YPW1784 to obtain a large number of colonies. After growth on selective medium (Synthetic Complete medium with 0.9 g/L 5-FluoroOrotic Acid), 244 colonies selected regardless of their fluorescence level were streaked on fresh plates (again SC + 5-FOA medium) and then replica plated on YPD + Hygromycin B (10 g/L Yeast extract, 20 g/L Peptone, 20 g/L Dextrose and 300 mg/L Hygromycin B) for negative screening. 106 of the resulting strains turned out not to be fluorescent, among which the vast majority were resistant to Hygromycin B, suggesting they were false positive transformants. The remaining 138 strains were all fluorescent and sensitive to Hygromycin B, as expected from true positive transformants. We then tried to amplify *P*_*TDH3*_ in all 244 strains using oligonucleotides 1891 and 1208 (Table S5) and we only observed a band of correct size after electrophoresis for the 138 fluorescent strains. After Sanger sequencing of the PCR products for the 138 positive strains, the type and frequency of mutations observed in the TATA box were found to be very close to expectation (Table S1). Average expression level and expression noise were measured for all 138 strains as described below. This set of alleles showed broad variation in average expression level (Figure 1 – figure supplement 1) and had a lower expression noise than TFBS mutations with comparable average expression levels. We selected seven TATA box variants (Table S2) with expression levels ranging from 20% to 75% of wild type to be included in the final set of 43 *P*_*TDH3*_ alleles. One of the random TATA box mutants contained a PCR-induced mutation in the GCR1.1 TFBS and was also included in the final set (Var23 in Table S1 and S2).

#### TATA box and TFBS mutants

To obtain variation in expression noise at expression levels below 20%, we combined mutations in TFBS with mutations in the TATA box in 12 additional *P*_*TDH3*_ alleles (Table S1). Two TATA box variants with 25% and 50% median fluorescence levels were each combined with six different TFBS variants for which median expression ranged from 4% to 45% relative to wild type. The 12 variants were created by PCR SOEing as described above for the double TFBS mutants, except in this case oligonucleotides 2425 and 2788 were used to amplify the upstream *P*_*TDH3*_ fragments and oligonucleotides 2787 and 104 were used to amplify the downstream fragments (Table S5). All 12 variants were transformed in strain YPW1784 and confirmed by Sanger sequencing.

#### Double-copy constructs

To create variation in average expression level and expression noise for expression levels more than 75% of wild type, we constructed 13 alleles with two copies of the whole *P*_*TDH3*_-*YFP-TCYC1* construct inserted in tandem at the *HO* locus (Table S1). One of these constructs carried two copies of the wild type *TDH3* promoter, while the others carried mutated versions of *P*_*TDH3*_. We reasoned that the presence of a second copy of the construct would lead to overexpression of YFP, as shown previously (Kafri *et al.* 2016), while differences in noise between the different alleles should be conserved. To construct these alleles, we first fused the selectable marker *URA3* upstream of the *P*_*TDH3*_-*YFP*-*T*_*CYC1*_ allele located at the right end of each of the final constructs (“*CONSTRUCT.2*” in Table S1) using PCR SOEing. *URA3* was amplified from the *pCORE-UH* plasmid using oligonucleotides 2688 and 2686 and the 13 *P*_*TDH3*_-*YFP*-*T*_*CYC1*_ constructs were amplified from the strains carrying the corresponding *P*_*TDH3*_ alleles using oligonucleotides 2687 and 1893 (Table S5). *URA3* and *P*_*TDH3*_-*YFP*-*T*_*CYC1*_ were then fused by overlap extension and the resulting fragments were amplified with oligonucleotides 2684 and 2683 (Table S5). Finally, each of the 13 different *URA3*-*P*_*TDH3*_-*YFP*-*T*_*CYC1*_ PCR products was transformed in the strain carrying the desired allele of *P_TDH3_-YFP-T_CYC1_* (strain carrying “*CONSTRUCT.1*” in Table S1). During these transformations, the *KanMX4* drug resistance cassette was replaced with *URA3-P_TDH3_-YFP-T_CYC1_* by homologous recombination so that the final constructs were *ho::P_TDH3_-YFP-T_CYC1_-URA3-P_TDH3_-YFP-T_CYC1_*. To control for the impact of the URA3 marker on the activity of the *TDH3* promoter, we constructed strain YPW2675 (*ho::P_TDH3_-YFP-T_CYC1_-URA3*) by replacing the *KanMX4* cassette with *URA3* amplified using primers 2684 and 2685 (Table S5).

YPW2675 was used as the reference when reporting the expression phenotypes (median and noise) of the alleles with two copies of *P_TDH3_-YFP-T_CYC1_*. To validate the sequence of the full (5.2 kb) constructs, we performed two overlapping PCRs using oligonucleotides 2480 and 1499, and 1872 and 2635 (Table S5). PCR products were sequenced using primers 2480, 1499, 1204, 1872, 2635, 2686, 1305 and 601 in Table S5) to confirm they contained the correct *P*_*TDH3*_ alleles. However, using this PCR approach, insertion of more than two tandem copies of *P_TDH3_-YFP-T_CYC1_* would remain undetected. Therefore, we used quantitative pyrosequencing to determine the exact number of copies inserted in the 13 strains. We took advantage of the fact that all *P*_*TDH3*_ alleles inserted at *HO* carried the mutation A293g upstream of the start codon, while the endogenous *TDH3* promoter did not. This allowed us to determine the total number of *P*_*TDH3*_ copies at the *HO* locus by quantifying the relative frequency of A and G nucleotides at position −293 across all copies of the *TDH3* promoter in the genome. For instance, if only one copy of *P*_*TDH3*_ is present at *HO*, then the frequency of G at position −293 is expected to be 0.5, since there is one copy of the G allele at the *HO* locus and one copy of the A allele at the endogenous *TDH3* locus. If two copies are present at HO, a frequency of 2/3 is expected for G, and if three copies are present, a frequency of 0.75 is expected. To determine these allele frequencies, we amplified *P*_*TDH3*_ in five replicates from all strains carrying two copies of the construct as well as from YPW2675 carrying a single copy using oligonucleotides 2268 and 3094 (Table S5). PCR products were denatured and purified using a PyroMark Q96 Vacuum Workstation (Qiagen) and pyrosequencing was performed on a PyroMark Q96 ID instrument using oligonucleotide 2270 for sequencing (Table S5). Allele frequencies were determined from the relative heights of the peaks corresponding to the A and G alleles on the pyrograms, with the typical correction factor of 0.86 applied to A peaks. Using this method, a small but significant bias toward the G allele was detected, as the observed frequency of G in strain YPW2675 was 0.55 instead of 0.5. This could be caused by PCR bias due to the fact that the A and G alleles are located at different genomic positions. We applied the linear correction y = x * (0.5/0.45) – 0.111 to remove the effect of this PCR bias when calculating the frequency of G alleles. Overall, we found that six strains had a frequency of G significantly higher than 2/3 (*t*-test, *P* < 0.05). This suggested that these strains carried more than two copies of the *P*_*TDH3*_-*YFP*-*T*_*CYC1*_ construct and they were therefore removed from all subsequent analyses (except Var42 for reasons explained below).

#### Extra mutations

Sanger sequencing revealed that a substantial fraction of all *P*_*TDH3*_ alleles constructed (~25% of sequenced strains) carried an indel of one nucleotide in one of the homopolymer runs present in the promoter (Table S1). These mutations probably result from polymerase slippage during PCR amplification. For some *P*_*TDH3*_ alleles, we were able to isolate independent clones that differed only by the presence or absence of these homopolymer mutations, giving us the opportunity to test the impact of homopolymer length variation on transcriptional activity. Using the fluorescence assay described below, we found that *del431A*, *del54T* and *ins432A* had no detectable effect on median expression level or expression noise (Figure 2 – figure supplement 2C,D). Therefore, strains carrying these mutations were included in the expression and fitness assays.

### Yeast strains: construction of strains used to assay fitness

The strains described above all carried the *ho*::*P*_*TDH3*_-*YFP*-*T*_*CYC1*_ reporter construct, allowing sensitive quantification of the transcriptional activity of different *P*_*TDH3*_ alleles. In these strains, the endogenous promoter driving expression of the native TDH3 protein was left unaltered. To measure how variation in TDH3 protein levels induced by mutations in the *TDH3* promoter could impact cell growth, we inserted the final set of 43 *P*_*TDH3*_ alleles described above upstream of the endogenous *TDH3* coding sequence (Table S1). *P*_*TDH3*_ variants were integrated in the genetic background of strain YPW1001, which is almost identical to the reference strain YPW1002 used for the expression assays, except that the mating type of YPW1001 is *MAT***a** and it carries a *P*_*TDH3*_-*YFP*-*T*_*CYC1*_-*NatMX4* construct at *HO* conferring resistance to Nourseothricin. The reporter construct served a dual purpose: it ensured that the strains used in the expression and fitness assays carried the same number of copies of *TDH3* promoter in their genomes and it allowed high-throughput counting of yellow-fluorescent cells carrying *P*_*TDH3*_ variants in the competition experiments described below. Importantly, we did not detect any difference in fluorescence levels between strains YPW1002 and YPW1001 (Figure 2 – figure supplement 2A,B), indicating that the few genetic differences between the background of the strains used in the expression and fitness assays did not significantly affect the activity of the *TDH3* promoter.

#### Single-copy constructs

To insert the 35 alleles containing a single copy of *P*_*TDH3*_ at the native *TDH3* locus, we first replaced the endogenous *TDH3* promoter of strain YPW1001 with a *CORE-UK* cassette (*URA3-KanMX4*) amplified with oligonucleotides 1909 and 1910 (Table S5) to create strain YPW1121. Then, the 35 *P*_*TDH3*_ alleles were amplified from the *HO* locus in the strains previously constructed (Table S1) using oligonucleotides 2425 and 1305 (Table S5). PCR products were purified using a DNA Clean & Concentrator kit (Zymo Research), amplified using primers 1914 and 1900 (Table S5) to attach appropriate homology tails and transformed in strain YPW1121. In addition, because all the *P*_*TDH3*_ variants inserted at *HO* carried the PCR-induced mutation A293g, we created the strain YPW1189 that carried mutation A293g in the endogenous *TDH3* promoter. YPW1189 served as the reference strain when calculating relative fitness. In all these strains, the presence of the correct mutations in *P*_*TDH3*_ at the native locus was confirmed by Sanger sequencing of PCR products obtained with oligonucleotides 1345 and 1342 (Table S5).

#### Double-copy constructs

To measure the impact on fitness of overexpression of the native TDH3 protein, we created 7 tandem duplications of the whole *TDH3* locus (*TDH3*::*P*_*TDH3*_-*TDH3*-*URA3*-*P_TDH3_*-*TDH3*) that contained the same combinations of promoter alleles as those inserted at *HO* (Table S1). Duplications of *TDH3* were built in a similar way as the double-copy constructs inserted at *HO*. First, *URA3* was amplified from the *pCORE-UH* plasmid using oligonucleotides 2688 and 2686 and the *TDH3* variants corresponding to the copy located on the right in the final constructs (“*CONSTRUCT.2*” in Table S1) were amplified using oligonucleotides 2687 and 1893 (Table S5). *URA3* and *P*_*TDH3*_-*TDH3* PCR products were then fused by overlap extension and the resulting fragments were amplified with oligonucleotides 2696 and 2693 (Table S5). Finally, each of the 7 different *URA3-P_TDH3_-TDH3* products was transformed in the strain carrying the desired allele for the left *P*_*TDH3*_-*TDH3* copy (“*CONSTRUCT.1*” in Table S1). To control for the impact of URA3 expression on fitness, we constructed strain YPW2682 (*TDH3::P_TDH3_-TDH3-URA3*) by transforming a *URA3* cassette amplified from plasmid *pCORE-UH* with oligonucleotides 2696 and 2697 in strain YPW1189. YPW2675 was used as the reference when reporting the relative fitness of the 7 strains carrying two copies of *TDH3*. To sequence the full *TDH3* duplications (5.5 kb), we performed four overlapping PCRs using oligonucleotides 1345 and 1499, 2694 and 1911, 2670 and 1342, 601 and 2695 and sequenced them with oligonucleotides 1345, 1499, 601, 2691, 2053, 2670, 1342, 601, 2695 (Table S5).

As described for the double-copy constructs at *HO*, we used quantitative pyrosequencing to determine the exact number of *TDH3* copies inserted in the 7 strains. However, we could not directly quantify the frequency of mutation A293g in these strains, because all copies of *TDH3* promoters present in their genomes carry the G mutation. Therefore, we first crossed all 7 strains to YPW1139 (Metzger *et al.* 2016), a strain that contains the A allele at position −293 of the native *TDH3* promoter. In the resulting diploids, the frequency of G allele at the native *TDH3* locus is expected to be 0.5 if the original haploid strain carried a single copy of *TDH3* at the native locus, 2/3 if it carried two copies of *TDH3* at the native locus and 3/4 if it carried three copies. To determine allele frequency at position −293 of *P*_*TDH3*_ for the native *TDH3* locus only, we amplified the promoter using primers 2268 and 3095 specific to the native locus (Table S5) and then used pyrosequencing as described above. We found that one strain carried three copies of *TDH3* at the native locus instead of two (Table S1). However, we did not exclude the corresponding variant (Var42) from subsequent analyses, because it also integrated three copies of the reporter construct at *HO*.

Finally, during growth rate assays, cells carrying a tandem duplication of *TDH3* could potentially lose a copy of *TDH3* through intrachromosomal homologous recombination, which could affect fitness estimates. In strains carrying *TDH3::TDH3-URA3-TDH3* constructs, the loss of a *TDH3* copy by recombination should be accompanied by the deletion of the *URA3* marker. To estimate how frequently such recombination events might occur, we quantified the frequency of Ura-cells in strain YPW2679 (*TDH3::TDH3-URA3-TDH3*) at four time points over the course of 50 generations of growth in similar conditions as used in competition growth assays. Four replicate cultures of YPW2679 were grown to saturation in SC - Ura medium at 30°C. Then, 0.1 ml of each culture was plated on SC + 5-FOA medium and each culture was diluted to a density of 10^4^ cells/ml in YPD rich medium. Dilution to 10^4^ cells/ml in YPD was repeated every 12 hours for 72 hours and plating on SC + 5-FOA was repeated every 24 hours. After three days of incubation at 30°C, colony-forming units were counted on all SC + 5-FOA plates, allowing the estimation of the frequency of Ura-cells every ~17 generations for a total of ~50 generations (Table S6). The frequency of Ura-cells was found to increase during the first 34 generations of growth before reaching a plateau representing a state of mutation-selection balance. At this stage, the average frequency of Ura-cells was about 5.2 x 10^−5^. Therefore, even if spontaneous loss of one *TDH3* copy occurred in a fraction of cells, these events were found to be too rare to have a significant impact on fitness estimates.

#### TDH3 deletion

We deleted the native *TDH3* locus in the genetic background of strain YPW1001 to create strain YPW1177. To do this, we amplified a region of 171 bp immediately upstream of the *TDH3* promoter using oligonucleotides 1345 and 1962 (Table S5). Oligonucleotide 1962 is composed of a 5’ sequence of 22 nucleotides priming directly upstream of the *TDH3* promoter fused to a 3’ sequence of 38 nucleotides homologous to the 3’UTR sequence immediately downstream of *TDH3* coding sequence. Therefore, transformation of the PCR product in strain YPW1121 (*tdh3::URA3-KanMX4-TDH3*) led to the deletion of the *URA3-KanMX4* cassette and of the *TDH3* coding sequence. In this strain, both the *TDH3* promoter and the *TDH3* coding sequence are deleted, and the coding sequence of the upstream gene *PDX1* is fused to the terminator sequence of *TDH3*, so that *PDX1* would remain functional. Correct deletion of *TDH3* was confirmed by Sanger sequencing of the region amplified with oligonucleotides 1345 and 2444 (Table S5) in strain YPW1177.

#### GFP competitor

To measure how variation in *TDH3* expression affected growth rate, the strains described above were all grown competitively against a common strain, YPW1160, which carried a *P*_*TDH3*_-*GFP*-*T*_*CYC1*_-*KanMX4* construct inserted at the *HO* locus in the same genetic background as the other strains. The expression of Green Fluorescent Protein in YPW1160 cells allowed for highly efficient discrimination from cells expressing the Yellow Fluorescent Protein using flow cytometry. To construct strain YPW1160, the *GFP-T_CYC1_* sequence was amplified from strain YPW3 (*swh1*::*P*_*TDH3*_-*GFP*-*T_CYC1_*, obtained from Barry Williams) using oligonucleotides 601 and 2049 (Table S5). In parallel, *KanMX4* was amplified from strain YPW1002 using oligonucleotides 2050 and 1890 (Table S5). The two fragments were fused by PCR SOEing and the product was amplified using oligonucleotides 601 and 1890 (Table S5) before transformation in strain YPW1001 (*ho*::*P*_*TDH3*_-*YFP*-*T*_*CYC1*_-*NatMX4*). Selection on G418 allowed the recovery of cells that switched the *YFP*-*T*_*CYC1*_-*NatMX4* cassette for the *GFP*-*T*_*CYC1*_-*KanMX4* cassette. The fluorescence emission detected on the flow cytometer was consistent with expression of GFP.

### Expression assays

#### Quantification of fluorescence using flow cytometry

Fluorescence level was measured as a proxy for *P*_*TDH3*_ transcriptional activity using flow cytometry as described in (Metzger *et al.* 2016). All strains were revived from −80°C glycerol stocks on YPG plates (10 g Yeast extract, 20 g Peptone, 30 ml Glycerol, 20 g agar per liter) and, after 2 days of growth, arrayed in 96-well plates containing 0.5ml of YPD medium per well. In addition to the tested strains, the reference strain YPW1002 was inoculated in 24 positions, which were used to correct for plate and position effects on fluorescence. Strain YPW978, which does not contain the *YFP* reporter construct (Metzger *et al.* 2016), was inoculated in one well per plate and used to correct for autofluorescence. Cells were maintained in suspension at 250 rpm by the presence of a 3-mm glass bead in each well. After 20 hours of growth at 30°C, cells were transferred on YPG omnitrays using a V&P Scientific pin tool and grown for 2 days. Next, samples from each omnitray were inoculated in six replicate 96-well plates in 0.5 ml of YPD and grown for 22 hours at 30°C until they reached saturation. At this point, 15 μl of each culture was diluted into 0.5 ml of PBS (phosphate-buffered saline) in 96-well plates. Fluorescence was recorded for ~20,000 events per well using a BD Accuri C6 instrument coupled with a HyperCyt autosampler (IntelliCyt Corp). A 488-nm laser was used for excitation and a 530/30 optical filter for acquisition of the YFP signal. A modified version of this protocol was used to measure the fluorescence of the final set of 43 *P*_*TDH3*_ variants with experimental conditions more similar to those experienced during the competition growth assays. After the 22 hours of growth in YPD, samples were not immediately run on the flow cytometer, but instead they were diluted to fresh medium every 12 hours for 36 hours to reach steady exponential phase of growth. Prior to each dilution, cell density was measured for all samples using a Sunrise absorbance reader (Tecan) and one dilution factor was calculated for each 96-well plate so that the average cell density would reach 5 x 10^6^ cells/ml after 12 hours of growth. This procedure ensured that all samples were maintained in constant exponential growth since no sample reached a density above 10^7^ cells/ml, while limiting the strength of genetic drift since the number of cells transferred during dilution was larger than 10,000. Another difference with the protocol mentioned above is that no glass bead was added to the plates. Instead, cells were maintained in suspension by fitting the culture plates on a rotating wheel. After 36 hours of growth, samples were diluted to 2.5 x 10^6^ cells/ml in PBS and the fluorescence of 20,000 events per well was acquired by flow cytometry. All flow data are available in the FlowRepository under Repository ID xxxxx.

#### Relationship between mRNA levels and fluorescence

The relationship between fluorescence intensity measured by flow cytometry and fluorophore concentration in a cell is expected to be positive and monotonic, but this relationship is not necessarily linear (Wang and Gaigalas 2011). For most flow cytometers, the photomultiplier tube (PMT) voltages can be calibrated to approach a linear relationship for the range of fluorescence intensities covered by the samples, but this cannot be done on the Accuri C6 because PMT voltages are fixed. Instead, we empirically determined the function relating fluorescence intensities to *YFP* mRNA levels using eight strains with different fluorescence levels and then we applied this function to transform fluorescence intensities for each cell of every sample. The function between fluorophore concentration (*y*) and fluorescence intensity (*x*) was previously determined to be of the form log(*y*) = a × log(*x*) + *b* (Wang and Gaigalas 2011). In our case, ! represents mRNA level instead of fluorophore concentration, but this should not affect the shape of the function since previous studies found a linear relationship between mRNA levels and fluorophore concentration (Wolf *et al.* 2015; Kafri *et al*. 2016). To determine the constants *a* and *b*, we measured fluorescence intensity and *YFP mRNA* levels in eight strains covering the whole range of fluorescence levels expressed by the strains included in this study. First, three replicates of YPW978 (non-fluorescent strain), YPW2683 (*ho*::*P*_*TDH3*_-*YFP*-*T*_*CYC1*_/*ho*::*P*_*TDH3*_-*GFP*-*T*_*CYC1*_ diploid) and seven strains carrying variants of the *ho*::*P*_*TDH3*_-*YFP*-*T_CYC1_* construct with different *P*_*TDH3*_ alleles (Table S1) were grown for 24 hours at 30°C in 5 ml of YPD, along with 24 replicates of strain YPW1182 expressing GFP (same genetic background as YPW1160, except with *MATα* mating type). All samples were then diluted to a density of 2 x 10^6^ cells/ml in fresh YPD medium and grown for another 4 hours. Next, 0.1 ml of each culture was transferred to 0.4 ml of PBS and fluorescence intensity was immediately scored for ~20,000 events per sample on the BD Accuri C6 instrument.

In parallel, for each replicate of the tested strains, 0.5 ml of culture was mixed with 0.5 ml of one of the 24 cultures of YPW1182 strain in a microcentrifuge tube. Genomic DNA and RNA were co-extracted from the 24 mixed populations using a modified version of Promega SV Total RNA Isolation System and cDNA was synthesized from RNA samples as previously described (Metzger *et al.* 2015). Then, pyrosequencing was performed to quantify the relative frequency of *YFP* and *GFP* sequences in gDNA and cDNA samples (Wittkopp 2011). The pyrosequencing assay was designed to quantify allele frequency at a position located 607 bp downstream of the *YFP* start codon, for which a GT/TA difference exists between *YFP* and *GFP* coding sequences. A region that encompassed this polymorphism was amplified in all gDNA and cDNA samples using oligonucleotides 2723 and 2725 (Table S5). These oligonucleotides were designed to anneal both to *YFP* and *GFP* coding sequences, which are 98% identical. Pyrosequencing was performed on a PyroMark Q96 ID instrument using oligonucleotide 2726 for sequencing (Table S5). Because the sequenced region contained two variable positions (G/T and T/A), we determined allele frequencies separately for each position and used the average as the relative frequency of *YFP* and *GFP* alleles. For each sample, we then calculated *YFP* mRNA level relative to the reference strain YPW1002 using the measured frequency of *YFP* allele in gDNA and cDNA. First, we corrected for small biases in allele frequencies that could be caused by PCR bias. To do this, we took advantage of the fact that true allele frequencies were known for gDNA samples of YPW1002 (100% *YFP*), YPW2683 (50% *YFP*) and YPW1182 (0% *YFP*). We regressed the measured allele frequencies on the true allele frequencies for all gDNA replicates of these three samples using R *smooth.spline* function. The fitted model was then used to correct allele frequencies for all other gDNA and cDNA samples. Our next goal was to calculate., defined as the abundance of *YFP* mRNA expressed by each tested strain relative to the abundance of *GFP* mRNA expressed by YPW1182. If / is the frequency of *YFP* allele in the gDNA sample and 0 the frequency of *YFP* allele in the cDNA sample, then 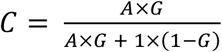. From this equation, we can deduce that 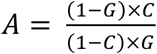. We applied this formula to our measured estimates of *G* and *C* to calculate *A*. For each sample, the calculated value of. was divided by the value obtained for the reference strain YPW1002 to obtain an estimate of *YFP* mRNA level expressed relative to the reference. Finally, we identified the function of form log(*y*) = *a* × log(*x*) + *b* that best fitted to our measures of mRNA levels and fluorescence intensities using R function *nls*. The least-square estimates of the parameters were a = 10.469 and b = −9.586. As expected, we observed a nonlinear relationship between *YFP* mRNA level and fluorescence intensity (Figure 1B, *R*^2^ = 0.83), but a linear relationship between the logarithm of *YFP* mRNA level and the logarithm of fluorescence intensity (Figure 1C, *R*^2^= 0.99).

#### Analysis of flow cytometry data for expression

Flow cytometry data were analyzed using R packages *flowCore* (Hahne *et al.* 2009) and *flowClust* (Lo *et al.* 2009) with modifications of the methods described in Duveau, Toubiana, *et al.* (2017) linked to the transformation of fluorescence intensities mentioned above. First, the clustering functions of *flowClust* were used to filter out all events that did not correspond to single cells based on the height and the area of the forward scatter signal. Then, the intensity of the fluorescence signal was scaled by cell size in several steps. We first performed a principal component analysis on the logarithm of forward scatter (FSC.A) and logarithm of fluorescence (FL1.A) for all filtered events. Next, we defined the vector *v* between the origin and the intersection of the two eigenvectors. We then calculated the angle *θ* between the first eigenvector and *v*. FSC.A and FL1.A data were transformed by a rotation of angle *θ* centered on the intersection between the two eigenvectors. Finally, for each event, the transformed FL1.A value was divided by the transformed FSC.A value to obtain a measure of fluorescence level independent of cell size. The fluorescence level of each individual cell was then rescaled using the function log(*y*) = 10.469 × log(*x*) − 9.586 to follow a linear relationship with *YFP* mRNA levels, as explained in the previous paragraph. For each sample, the median *m*_*YFP*_ and the standard deviation *s*_*YFP*_ of expression were calculated from the fluorescence levels of at least 1000 cells. Next, we corrected for variation in fluorescence levels caused by factors beyond experimental control by using the 24 control samples present on each plate at the same positions. For each environment, log(*m*_*YFP*_) and log*s*_*YFP*_/log(*m*_*YFP*_) of control samples were fitted to a linear model that included explanatory variables such as average cell size, replicate, plate, row, column and flow run. The variable that had the greatest impact on fluorescence was found to be “flow run”. This effect is caused by random variation in the sensitivity of the flow cytometer to measure fluorescence intensity between each run of 48 samples, rather than actual variation in YFP expression, as indicated by the observation of random shifts when running the same plate multiple times. Therefore, for each sample, the effect of “flow run” was removed on a scale that was linearly related to fluorescence intensity and not mRNA level. Given that the logarithm of mRNA levels log(*y*) scales linearly with the logarithm of fluorescence intensity, the linear model to correct for “flow run” and “row” effects was applied on linear estimates of *median*(log(*y*)) and *σ*(log(*y*)). log*m*_*YFP*_ scales linearly *median*(log(*y*)) and log(*s*_*YFP*_)/log(*s*_*YFP*_) is expected to scale approximately linearly with *σ*(log(*y*)). Indeed, the delta method (Ver Hoef 2012) postulates that *θ*(*f*(*x*)) = *σ*^2^ × (*f*′(*x*))^2^ and the first derivative of log(*x*) is 1/*x*. The corrected values of log(*m*_*YFP*_) and log(*s*_*YFP*_)/log(*m*_*YFP*_) were then used to calculate corrected values for log(*m*_*YFP*_) and log(*s*_*YFP*_). This procedure was found to uniformly decrease the variance of *m*_*YFP*_ and *s*_*YFP*_ among replicates of a same strain, independently of the average *m*_*YFP*_ of the strain. Next, we corrected for autofluorescence by subtracting the mean values of *m*_*YFP*_ and *s*_*YFP*_ among replicates of the non-fluorescent strain YPW978 from the values of *m*_*YFP*_ and *s*_*YFP*_ of each sample. At this stage, in addition to *s*_*YFP*_, we calculated three other metrics of expression variability (i.e., noise), *CV\** = *s*_*YFP*_/*m*_*YFP*_, *logCV*\* = log_10_(*s*_*YFP*_/*m*_*YFP*_) and *Noise strength* = *s*_*YFP*_^2^/*m*_*YFP*_. These metrics are similar to the coefficient of variation, the logarithm of the coefficient of variation and the Fano factor (Kaern *et al.* 2005), except that *m*_*YFP*_ is a median instead of a mean. The four metrics of expression noise were used in parallel in all subsequent analyses. For each strain, samples for which *m*_*YFP*_ and *s*_*YFP*_ departed from the median value among replicate populations by more than four times the median absolute deviation were discarded. For each sample, median expression and expression noise were then divided by the mean phenotypic value obtained among replicate populations of the reference strain YPW1002 (for single-copy *P*_*TDH3*_ variants) or YPW2675 (for double-copy *P*_*TDH3*_ variants). Finally, these relative measures of median expression and expression noise were averaged among replicate populations of each genotype.

### Fitness assays

#### Doubling time in each environment

Prior to performing competition assays, we measured the doubling time of the reference strain YPW1160 when grown in YPD medium. Three replicate cultures of YPW1160 were started in parallel in 5 ml of YPD and incubated for 36 hours at 30°C with dilution to 5 x 10^5^ cells/ml every 12 hours. After the last dilution, cell density T was quantified every 60 minutes for 10 hours and then after another 800 minutes by measuring optical density at 660 nm. Doubling time was calculated as the inverse of the slope of the linear regression of log(D) / log(2) on time during the logarithmic phase of growth where the relationship between log(D) and time is linear. The average doubling time of the reference strain YPW1160 used in subsequent competition assays was found to be 80 minutes in YPD.

#### Competition assays against a common reference

Because the deletion of *TDH3* is known to cause only a ~5% reduction in growth rate, detecting a significant impact on fitness of a change in *TDH3* expression level or expression noise required highly accurate measurements of growth rate. For this reason, we decided to use head-to-head competition assays between strains expressing different levels of TDH3 protein and a common reference strain (YPW1160) to measure relative growth rate, which is a more sensitive method than directly measuring the absolute growth rate of each isolated strain (Gallet *et al.* 2012). Indeed, the additive effect of micro-environmental variation on growth estimates is nullified during competitive growth, because the two competitors are grown in the same microenvironment. Relative growth rate during log-phase was used as a proxy for fitness in this study, although this is not the only component of fitness.

To identify experimental conditions that would allow accurate estimates of fitness (precision of at least 10^−3^) while keeping cost and labor reasonable, we first performed a power analysis based on simulations to determine how experimental parameters affected accuracy. We decided that during the competition assays cells would be maintained in the logarithmic phase of growth by repeated dilutions in small volume of medium in 96-well plates to handle large number of samples in parallel. Six different parameters associated with this experimental design were varied in the simulations: 1) The number of biological replicates for each sample; 2) The starting frequency of the two strains competed against each other; 3) The difference in fitness between the two competitors; 4) The number of generations after dilution to fresh medium, a parameter that determined the number of cells transferred (or the bottleneck size) after each dilution; 5) The number of dilution cycles, which also determined the number of times the relative frequency of the competed strains was assessed; 6) The number of cells counted each time the relative frequency of the competed strains was assessed. For each of 20,160 combinations of parameter values, the competition assay was simulated 5,000 times to estimate the standard deviation of the selection coefficient. Then, given this standard deviation and the tested number of replicates, R function *power.t.test* was used to determine the minimum difference in selection coefficient that could be detected with a significance level of 0.05 and a power of 0.95. All six parameters were found to have an impact on the precision of the selection coefficient, but to different degrees. Interestingly, precision was maximized for intermediate values of the number of generations between two consecutive dilutions and for intermediate values of the total number of dilution cycles, because of the impact of these parameters on genetic drift. To achieve a precision close to 10^−3^ in the actual competition experiment, we decided to use eight replicates per sample, to mix the two competing strains in equal proportion, to use a common competitor strain (YPW1160) that had a similar fitness as the wild-type strain (YPW1189), to grow cells for about eight generations of the common competitor after each dilution, to use a total of four phases of growth followed by dilutions and to score genotype frequencies at four time points by screening at least 50,000 cells per sample on the flow cytometer.

The tested strains carrying different alleles of the *TDH3* promoter at the native locus and expressing YFP (Table S2) were arrayed on four 96-well plates in 0.5 ml of YPD, with two replicates of each strain on each plate. In parallel, the common competitor YPW1160 expressing GFP was also arrayed on four 96-well plates in 0.5 ml of YPD. All plates were incubated for 24 hours at 30°C on a rotating wheel. After measuring cell densities using a Sunrise plate reader, an equal volume of YFP and GFP cell cultures were mixed together and diluted in 0.5 ml of YPD in four 96-well plates. The dilution factor was calculated for each plate based on the doubling time of the GFP strain (YPW1160) so that the average cell density would reach ~5 x 10^6^ cells/ml after 12 hours of growth. This procedure of cell density measurement and dilution followed by 12 hours of growth was repeated three times and constituted the acclimation phase of the experiment, during which the relative frequency of YFP and GFP strains was not recorded. After this acclimation phase, samples were diluted every 10 hours in fresh YPD for a total of 30 hours of exponential growth. Cell density was measured for all samples prior to each dilution. Immediately after dilution to fresh medium, samples were diluted in 0.3 ml of PBS to a final density of 1.5 x 10^6^ cells/ml in four 96-well plates and placed on ice to stop growth. ~75,000 events were recorded for each sample on a BD Accuri C6 flow cytometer, using a 488 nm laser for excitation and two different optical filters (510/10 and 585/40) to acquire fluorescence. These filters allowed separation of the GFP and YFP signals. With this protocol, the relative frequency of YFP and GFP cells was measured at four time points during the competition assays.

#### Analysis of flow cytometry data for fitness

The number of cells expressing either YFP or GFP was counted for each sample using custom R scripts. After log^10^ transformation of the raw data, artifacts were removed by excluding events with extreme values of forward scatter (FSC.A and FSC.H) or fluorescence intensity (FL1.H and FL2.H). FL1.H corresponds to the height of the fluorescence signal acquired through the 510/10 filter, which is more sensitive to GFP emission, and FL2.H corresponds to the height of the fluorescence signal acquired through the 585/40 filter, which is more sensitive to YFP emission. Next, a principal component analysis was performed on the logarithm of FL1.H and FL2.H. The first principal component captured differences in fluorescence caused by variation in cell size, while the second component captured differences in fluorescence between cells expressing YFP and GFP. We then computed the Kernel density estimate of the second component, which allowed the separation of three populations of cells: 1) GFP cells with high scores on the second component, 2) YFP cells with low scores and 3) a smaller population with intermediate scores considered as doublets, *i.e.* events corresponding to two cells scored simultaneously, one expressing YFP and the other GFP. Doublets for which the two cells expressed the same fluorophore should also occur at low frequency in the GFP and YFP populations, but these doublets cannot be distinguished from singletons based on fluorescence. The number of YFP cells *N*_*Y*_ and the number of GFP cells *N*_*G*_ can be calculated from the total number of YFP events *T*_*Y*_, the total number of GFP events *N*_*G*_, the number of YFP doublets *D*_*Y*_, the number of GFP doublets *D*_*G*_ and the number of YFP-GFP doublets *D*_*YG*_ using the following equations:

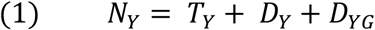

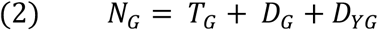

By analogy with the Hardy-Weinberg principle, we could expect that:

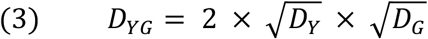

Therefore,

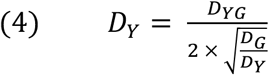

If we assume that doublets were formed randomly, then we should expect the same proportion of doublets in the YFP and GFP populations:

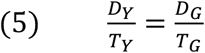

We can deduce from equations (1), (4) and (5) that:

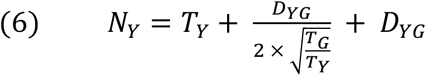

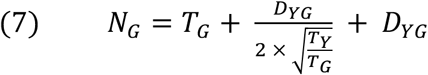

As all variables in the right-hand sides were known, we used equations (6) and (7) to estimate the number of YFP cells and the number of GFP cells in the sample. Then, for each sample, we determined the number of cell generations that occurred during the three dilution cycles, by using the measured cell densities before each dilution as well as the dilution factor. The median number of generations for all samples grown on a same 96-well plate was used as a rough estimate of the number of generations for the samples of the plate. The number of generations over the entire experiment was found to be about 22. The fitness of the YFP strain relative to the GFP competitor was calculated as the exponential of the slope of log_*e*_(*N*_*Y*_/*N*_*G*_) regressed on the number of generations at the four time points when genotype frequency was measured, based on equation (5.3) in (Cormack *et al.* 1990). For each tested strain, samples for which fitness departed from the median fitness among all eight replicate populations by more than four times the median absolute deviation were considered outliers and were excluded from further analysis. Outliers could occur for several reasons, one of them being the random appearance of a beneficial (or compensatory) *de novo* mutation during competitive growth (Gallet *et al.* 2012). For each sample, the fitness relative to the GFP strain was then divided by the mean fitness obtained for all replicate populations of the reference strain YPW1189 (for single-copy *P*_*TDH3*_ variants) or YPW2682 (for double-copy *P*_*TDH3*_ variants). We then calculated the mean relative fitness and standard deviation over the eight replicate populations of each tested strain. This measure of fitness expressed relative to a strain with the reference *TDH3* promoter sequence was used in all subsequent analyses.

#### Pairwise competition assays

To directly determine how expression noise impacts fitness at different levels of *TDH3* expression, we competed five pairs of strains with similar average expression levels but differences in *TDH3* expression noise, with each pair having a different average expression level (Table S3). This experimental design allowed differences in fitness caused by variation in noise to be directly observed without the assumption of transitivity (Gallet *et al.* 2012) and without the need to correct for the correlation between median expression and noise. In these experiments, we doubled the number of replicate populations (16) and the number of generations of growth (~42) to achieve greater precision in the fitness estimates for each pair of strains. We also measured the relative frequency of the two competitors using quantitative pyrosequencing instead of flow cytometry. This method did not require the expression of different fluorescent markers to distinguish cells from the two strains, allowing us to compete strains that only differed genetically by the mutations in the *TDH3* promoter affecting expression noise.

The competition assays were performed as described above, with the following differences in the protocol. First, strains with low noise and strains with high noise were arrayed each on two 96-well plates, with 16 replicates per genotype. After incubation on YPG omnitrays and growth to saturation in YPD, equal volumes of cultures of strains with low noise and high noise were mixed together and diluted in 0.5 ml of YPD. Following 36 hours of acclimation (as described above), six cycles of dilution followed by 12 hours of growth were performed. Dilution factors were calculated so that the average cell density on each plate would reach ~5 x 10^6^ cells/ml at the next dilution time point. Once every two cycles, the remaining cell cultures were centrifuged immediately after dilution and the pellets were stored in 30 μl of water at −80°C for later PCR amplification and pyrosequencing, so that genotype frequencies were quantified at four time points during the experiment. Cell pellets were thawed in the week after freezing and 15 μl of each sample was transferred in 30 μl of Zymolyase 20T (3 mg/ml) in 0.1M Sorbitol. After 15 minutes of incubation at 37°C, plates were vortexed vigorously for 15 seconds to disrupt cell wall and centrifuged at 3220 rcf for four minutes in an Eppendorf 5810 R centrifuge. 5 μl of supernatant were used as PCR template in 50 μl reactions that also included 1 μl of dNTPs (10 mM of each dNTP), 2.5 μl of forward and reverse primers (10 μM each), 10 μl of 5x KAPA2G Buffer B, 0.4 μl of KAPA2G Robust HotStart DNA polymerase (5U/μl) and 28.6 μl of water. The oligonucleotides used for each sample were specific to the target mutation in *P*_*TDH3*_ (Table S5). After 42 cycles of amplification, PCR products were denatured and purified using a PyroMark Q96 Vacuum Workstation (Qiagen) and pyrosequencing was performed on a PyroMark Q96 ID instrument using sequencing primers specific to the target mutations (Table S3 and S5). The frequency of the two genotypes was determined for each sample from the relative heights of the peaks corresponding to the two alternative nucleotides on the pyrograms. Samples with an average peak height below 5 were excluded, as this could result from weak PCR and cause biases in measured allele frequency. The number of generations between each time point was estimated using the cell densities before each dilution and the dilution factors as described above. Relative fitness was calculated as the exponential of the slope of log_*e*_(*f*_*H*_/*f*_*L*_) regressed on the number of generations across the four time points, where *f*_*H*_ and *f*_*L*_ were the relative frequency of the genotypes conferring high and low noise respectively (*f*_*H*_ + *f*_*L*_ = 1). Therefore, a fitness value above 1 meant that the strain with high noise grew faster than the strains with low noise, while a fitness value below 1 meant that the strain with low noise grew faster than the strain with high noise. For each pair of strains, replicates for which fitness departed from the median fitness across all 16 replicates by more than four times the median absolute deviation were considered outliers and were excluded.

### Analyzing the relationship between expression and fitness

The relationship between the average expression level of *TDH3* and fitness is not expected to follow a simple mathematical function. Therefore, we used LOESS regression to describe the relationship between median expression and fitness from the data collected with the set of 43 *P*_*TDH3*_ alleles, using the R function *loess* with a span of 2/3. Next, we tested the impact of expression noise on fitness, which was complicated by the fact that expression noise is correlated with median expression and by the fact that median expression is expected to have a larger impact on fitness than expression noise. To disentangle the effects of median expression and noise on fitness, we first calculated the residuals (ΔNoise) from a LOESS regression (span = 2/3) of expression noise on median expression. Next, we used a similar approach to calculate the residuals (ΔFitness) from a LOESS regression (span = 2/3) of fitness on median expression. ΔFitness is the variation in fitness that cannot be explained by a difference in median expression in our dataset. To test whether ΔFitness could be at least partially explained by variation in expression noise, we calculated the Pearson’s correlation coefficient between ΔNoise and ΔFitness and used the R function *cor.test* to test for significance of this correlation. We excluded the two strains that showed a median expression level above 125%, because the number of samples with high expression was too low for meaningful interpretation of ΔNoise and ΔFitness in this range of expression levels. In addition, we compared the correlations between ΔNoise and ΔFitness for two different classes of promoter variants determined based on their expression levels. First, we determined the maximum fitness from the LOESS regression of fitness on median expression. Next, we estimated the median expression value that would lead to a 0.005 reduction in fitness relative to the maximum. This expression value was used as a threshold to determine which strains had an expression close to the optimum or far from it. Three quantitative parameters were determined arbitrarily in these analyses: the span of the two LOESS regressions and the reduction in fitness used to determine the expression threshold. To test the robustness of the results to variation in these parameters, we calculated the correlations between ΔNoise and ΔFitness for 100 combinations of parameters where the span of the LOESS regressions took one of five values (2/6, 3/6, 4/6, 5/6 and 1) and the reduction in fitness took one of four values (0.0025, 0.005, 0.0075 and 0.01). All analyses were repeated for the four different metrics of noise mentioned above (Noise strength, SD, CV* and log(CV*)).

### Expression and fitness measured using TDH3-YFP fusion proteins

One important assumption in our analyses of the relationship between *TDH3* expression and fitness is that the median and noise of expression measured using the fluorescent reporter constructs inserted at *HO* are representative of the expression level of the TDH3 protein when the promoter variants are introduced at the native *TDH3* locus. To test whether the effects of mutations in the *TDH3* promoter were the same when introduced at *HO* or at the native *TDH3* locus, we constructed a *TDH3-YFP* fusion gene at the *TDH3* locus and then introduced 20 different *P*_*TDH3*_ alleles upstream of this reporter gene, including eight TFBS and four TATA box variants that were present in the competition assays (Table S1). To fuse the coding sequences of *TDH3* and *YFP*, we amplified the *YFP-T_CYC1_-KanMX4* construct from strain YPW1002 using primers 3415 and 3416 and transformed the PCR product in the non-fluorescent strain YPW978. Primer 3415 was designed to remove the stop codon of *TDH3* and the start codon of *YFP* and to insert a 30 bp linker between the coding sequences of the two genes (Huh *et al.* 2003). Then, the *TDH3* promoter was replaced with a *CORE-UH* cassette (*URA3-HphMX4*) amplified with oligonucleotides 1909 and 1910 (Table S5) to create strain YPW1618. The 20 *P*_*TDH3*_ alleles were amplified from the native locus in the strains previously constructed (Table S1) using oligonucleotides 1344 and 1342 (Table S5) and transformed into YPW1618 to replace the *CORE-UH* cassette. The presence of the expected mutations was confirmed by sequencing PCR products obtained with primers 1345 and 1952 (Table S5). The fluorescence level of the strains expressing the fusion proteins was measured in parallel to the fluorescence of strains carrying the same *P*_*TDH3*_ alleles at the *HO* locus. Four replicate samples of each strain were analyzed by flow cytometry after growth in YPD medium as described above. The expression of the reporter gene at the *HO* locus was found to be a strong predictor of the expression of the gene fusion at the native *TDH3* locus, both for median expression level (Figure 2 – figure supplement 1A, R^2^ = 0.99) and for expression noise (Figure 2 – figure supplement 1B, R^2^ = 0.76). These fusion protein alleles were not used to measure fitness effects of changing *TDH3* expression because the fusion protein caused a 2.5% reduction in fitness on its own, suggesting it altered TDH3 function. In addition, the impact of fusing YFP to TDH3 on fitness was quantified by comparing the competitive growth rate of strain YPW1002 expressing YFP from the *HO* locus to the growth rate of strain YPW1964 expressing the TDH3-YFP protein fusion. The expression of the fusion protein was found to cause a 2.5% reduction in fitness (Figure 2 – figure supplement 1C), which could be caused by altered function of the TDH3 protein when fused with YFP. For this reason, we decided not to use protein fusions to measure the fitness associated with different levels of TDH3 expression.

### Modeling the relationship between single cell expression level and population fitness

To understand how cell-to-cell variability in gene expression level could contribute to population fitness, we performed individual-based stochastic simulations of the growth of clonal populations of cells covering a wide range of mean expression and expression noise values of a single gene. All simulations were run as short experiments of fixed duration (1,000 minutes) where variability in expression level impacting single cell division rate was the only determinant of population growth rate. The behavior of the population was determined by: a) a normal distribution 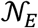 of expression levels for the focal genotype described by its mean *μ*_*E*_ and variance *σ*_*E*_^2^, and b) a function *DT* = *f*(*E*) relating single cell expression level *E* to the time in minutes separating two consecutive cell divisions, or doubling time *DT*. Single cell expression levels sampled from the expression distribution defined the doubling time for a given cell. Two different functions relating expression level to *DT* were explored: 1) a linear function (*DT* = −40 × *E* + 160) and 2) an inverted Gaussian function (*DT* = −160 × exp(− (*E* − 1)^2^/0.18) + 240). In each run of the simulation, a population of cells was tracked by recording information on the current expression level of each cell, the current *DT* derived from that expression level, and the amount of time remaining before the end of the experiment. For simplicity, the expression level of each mother and daughter cell was drawn from the normal distribution 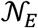 at each cell division and this expression level directly determined the *DT* value for the cell. To seed a starting population, 10^3^ cells were sampled from the expression distribution and their expression level was transformed into *DT*. To desynchronize the founding population, the initial values of *DT* were scaled by a random value between 0 and 1 to randomize the time to first division and a complete simulation was run. 10^3^ cells were drawn randomly at the end of the seed experiment and used to found a population for which growth rate was quantified. In the body of the simulation, each single cell was evaluated to determine if the current *DT* was greater than the remaining time in the experiment assessed for that cell, and if so, the cell divided, at which point new expression levels were drawn randomly from the normal distribution 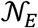 and independently for the mother and daughter cells. After cell division, the time remaining in the experiment for both mother and daughter cells decreased by the amount of the last *DT*, the new expression levels were translated into new values of *DT*, and the process repeated until *DT* values for all cells were greater than the remaining time in the experiment. Competitive fitness was calculated from the ratio of total number of cells *N*_*i*_ at the end of the experiment and the total number of cells *N*_*ref*_ obtained from simulating the growth of a reference genotype with mean expression *μ*_*ref*_ = 1 and noise *v*_*ref*_ = 0.1, as follows:

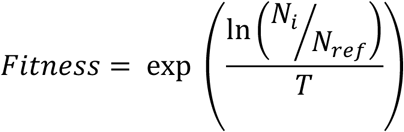

Mean expression *μ*_*E*_ of experimental genotypes were explored in the interval [0,2]. Noise values *v*_*E*_ were explored in the interval [0, 3] where noise was specified separately as standard deviation, coefficient of variation, and Fano factor. Experiment duration *T* was set at 1000 minutes for ease of computation. 100 replicates of each stochastic simulation were run to estimate 95% confidence intervals on fitness estimates. Simulations were coded in MATLAB R2015.

## Acknowledgements

We thank all members of the Wittkopp lab for helpful comments on the manuscript. This work was supported by a European Molecular Biology Organization postdoctoral fellowship (EMBO ALTF 1114-2012) to F.D., University of Michigan Rackham Graduate School (B.P.H.M.), National Institutes of Health Genome Sciences training grant (T32 HG000040 to B.P.H.M.), National Institutes of Health National Research Service Award (1F32GM115198) to A.H-D, and grants from the National Science Foundation (MCB-1021398) and National Institutes of Health (R01GM108826 and 1R35GM118073) to P.J.W.

## Supplementary figures

**Figure 1 – figure supplement 1.**
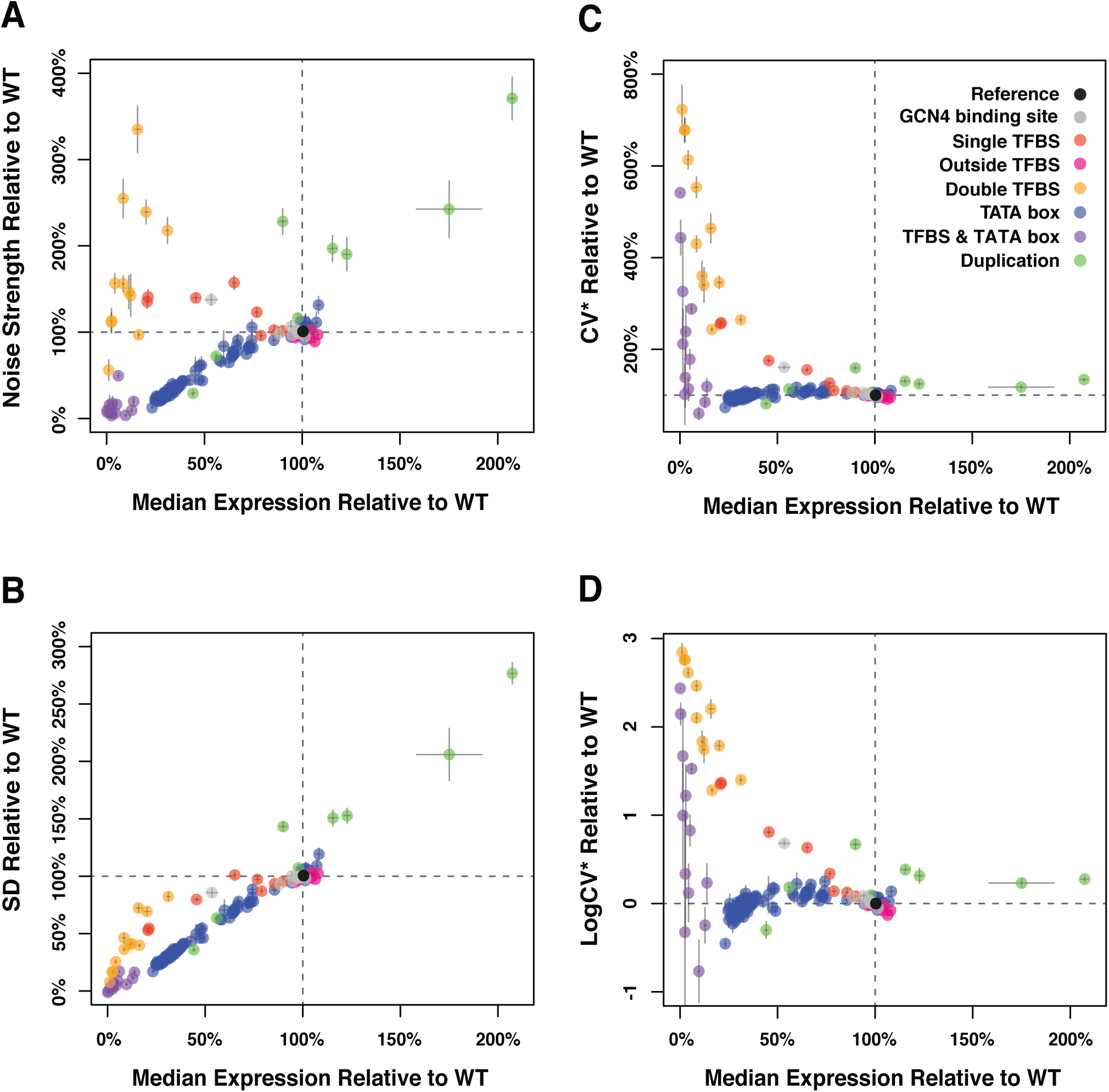
Median expression level and expression noise conferred by 171 variants of the *TDH3* promoter using four different metrics of noise. The four measures of expression noise are: **(A)** Noise strength, the variance divided by median fluorescence as in Figure 1 **(B)** SD, the standard deviation of fluorescence level among cells sharing the same genotype, **(C)** CV*, the standard deviation divided by median fluorescence level, and **(D)** LogCV*, the binary logarithm of CV*. Colors represent different categories of promoter variants based on the type of mutations they carry as indicated in **(B)**. The dotted lines show the activity of the wild type *TDH3* promoter. Error bars are 95% confidence intervals calculated from 6 replicates of each genotype.

**Figure 2 – figure supplement 1.**
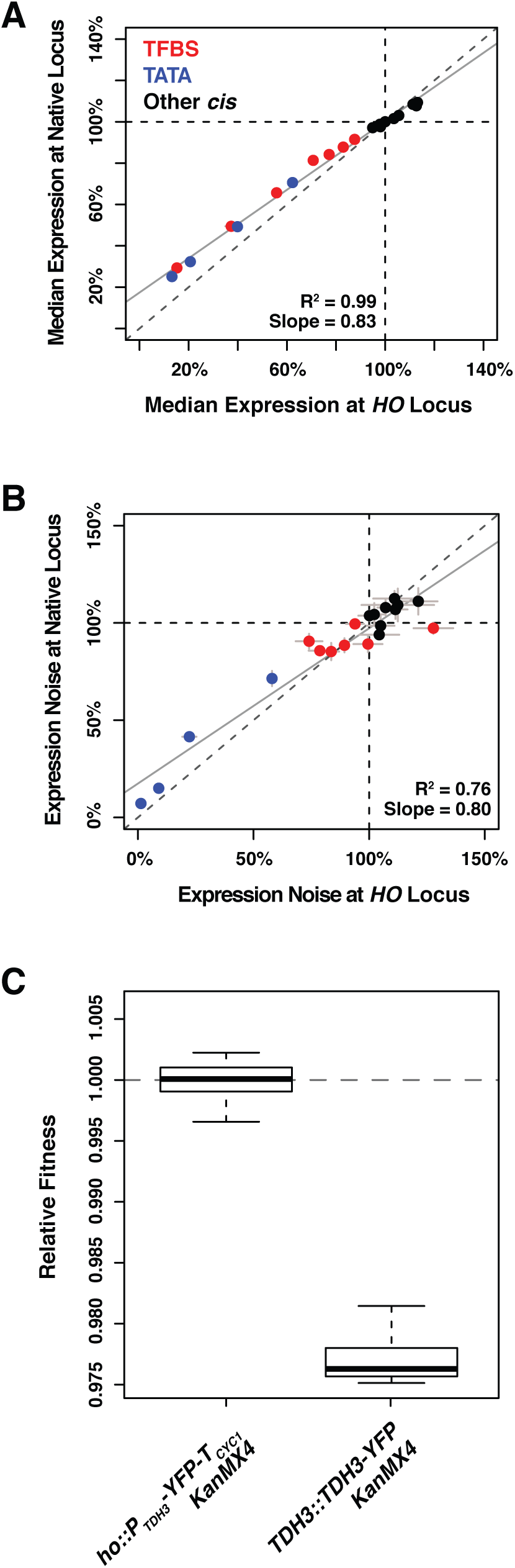
Comparing effects of 20 alleles of the *TDH3* promoter on expression of the YFP reporter at *HO* and of expression of the TDH3-YFP fusion at the native *TDH3* locus. **(A)** Median expression and **(B)** expression noise (noise strength) were quantified in six replicates for 20 pairs of strains carrying 20 different *TDH3* promoter variants inserted upstream of the *YFP* coding sequence at the *HO* locus or upstream of a *TDH3-YFP* gene fusion at the native *TDH3* locus. Colors represent different categories of promoter variants as shown in **(A)**. The dotted lines show the activity of the wild type *TDH3* promoter and the plain lines represent least square linear fits to data. Error bars are 95% confidence intervals. **(C)** Fitness consequence of the fusion of the YFP fluorescent protein at the C-terminus end of TDH3. Protein fusion was achieved with the strategy described in Huh *et al.* (2003), using the same polypeptide spacer of 10 amino acids between the two proteins. Fitness was measured for 12 replicate populations of two different genotypes. The first genotype (left) expresses *YFP* and *TDH3* under control of two separate copies of the *TDH3* promoter. The second genotype (right) expresses the *TDH3-YFP* gene fusion under control of the *TDH3* promoter, which caused a significant fitness reduction.

**Figure 2 – figure supplement 2.**
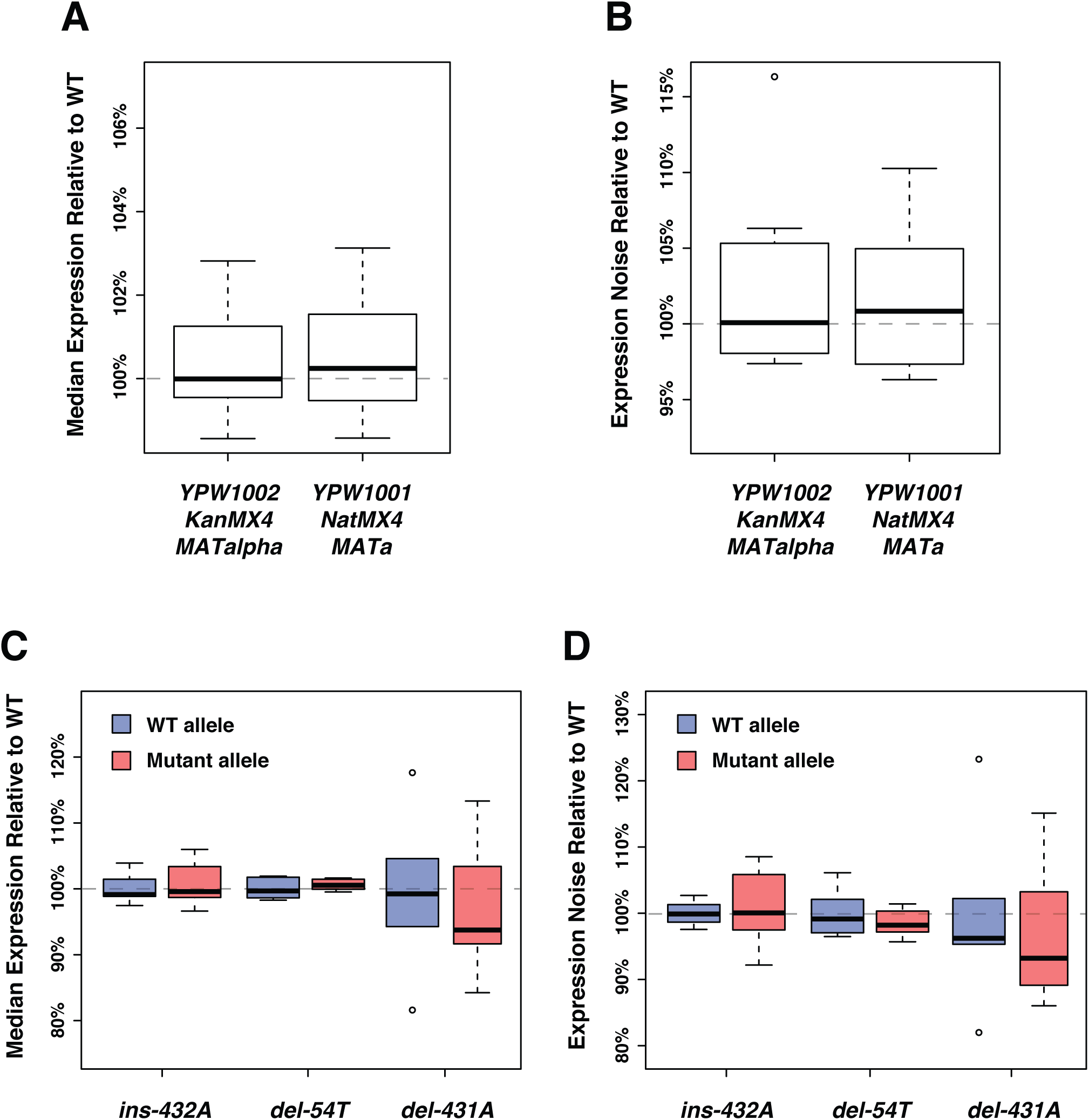
No significant impact of the genetic background on the expression of the fluorescent reporter. **(A)** Differences in drug resistance marker or mating type that exist between the strains used in the expression assays and the strains used in the fitness assays do not affect median expression. **(B)** The genetic changes mentioned in **(A)** do not significantly affect expression noise (noise strength). **(C)** No significant impact on median expression of three indels that frequently occurred during the construction of the different *P*_*TDH3*_ alleles. Position of each mutation is relative to the start codon. **(D)** The three indels mentioned in **(C)** do not significantly affect expression noise (noise strength). **(A-D)** Thick bars represent the median across six replicates. The bottom and top lines of each box represent 25^th^ and 75^th^ percentiles.

**Figure 3 – figure supplement 1.**
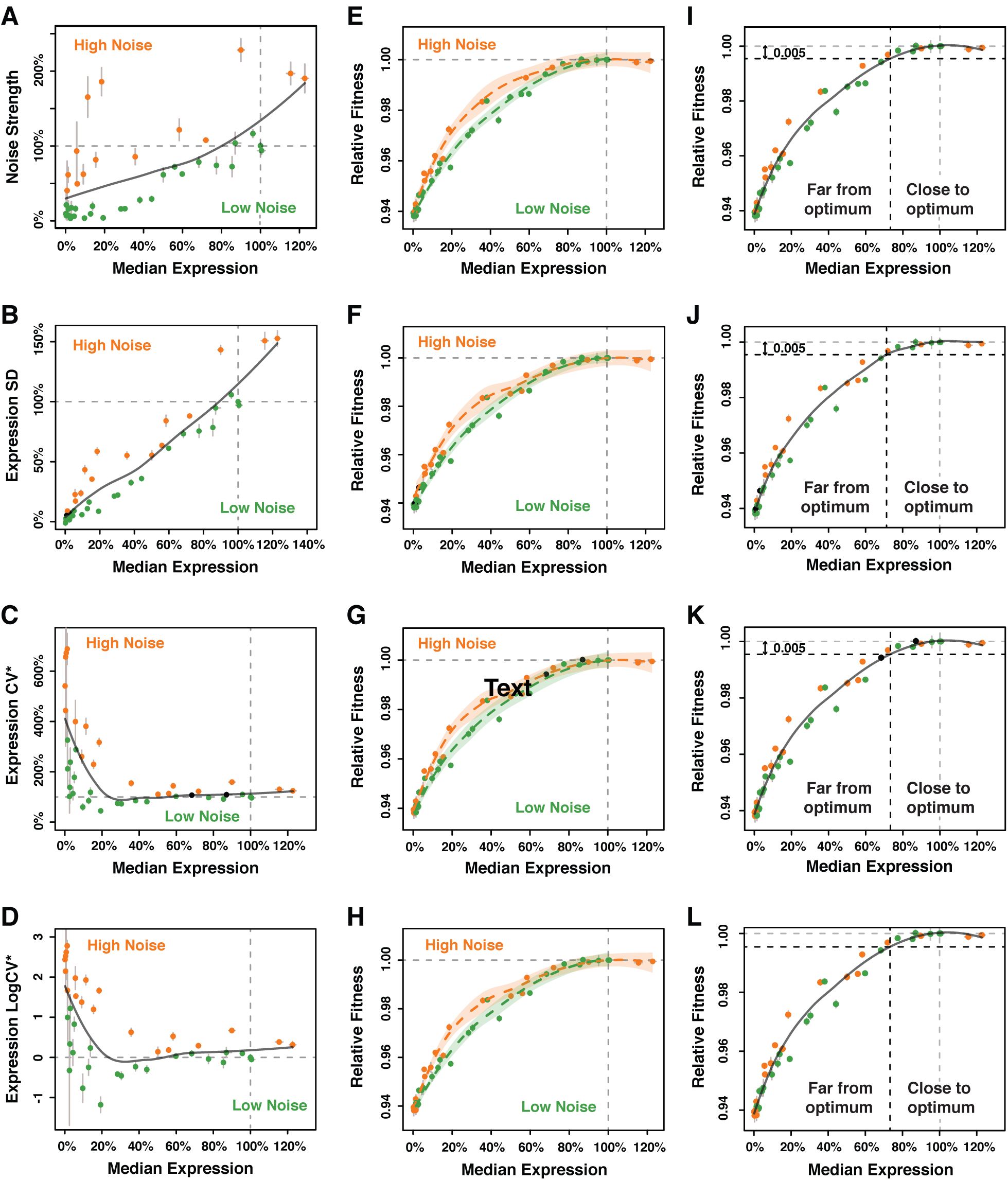
Calculation of ΔNoise and ΔFitness using four different metrics of noise. The four metrics of noise were: **(A,E,I)** the variance of expression divided by the median expression (noise strength), **(B,F,J)** the standard deviation of expression among genetically identical cells (SD), **(C,G,K)** the standard deviation divided by the median expression (CV*) and **(D,H,L)** the binary logarithm of the standard deviation divided by the median expression (LogCV*). **(A-D)** Separation of the 43 *P*_*TDH3*_ alleles in two categories based on their effects on median expression level and each of the four metrics of expression noise. The gray curve shows the LOESS regression of noise on median expression using a value of 2/3 for the smoothing parameter. Data points falling below the curve (green) correspond to *P*_*TDH3*_ alleles with low noise given their median level of activity. Data points above the curve (orange) correspond to *P*_*TDH3*_ alleles with high noise given their median activity. The residual of the LOESS regression (“ΔNoise”) is a measure of noise independent of median expression. **(E-H)** Distinct relationship between median expression level and fitness for strains with low noise (blue, ΔNoise* < −1%) and high noise (red, ΔNoise* > +1%). The two LOESS regressions were performed with smoothing parameter *α* equal to 2/3. **(I-L)** Partition of *P*_*TDH3*_ alleles in two groups based on the distance of their median activity to the optimal level of *TDH3* expression. The expression optimum (vertical gray dotted line) corresponds to the expression level predicted to maximize fitness from the LOESS regression of fitness on median expression (gray curve). The expression level at which the predicted fitness is 0.005 below the maximal fitness was chosen as the threshold (vertical black dotted line) separating promoters with median activity “close to optimum” from promoters with median activity “far from optimum”. The residual of the LOESS regression (“ΔFitness”) is a measure of fitness independent of the median *TDH3* expression level. **(A-L)** Error bars show 95% confidence intervals calculated from at least four replicate populations.

**Figure 3 – figure supplement 2.**
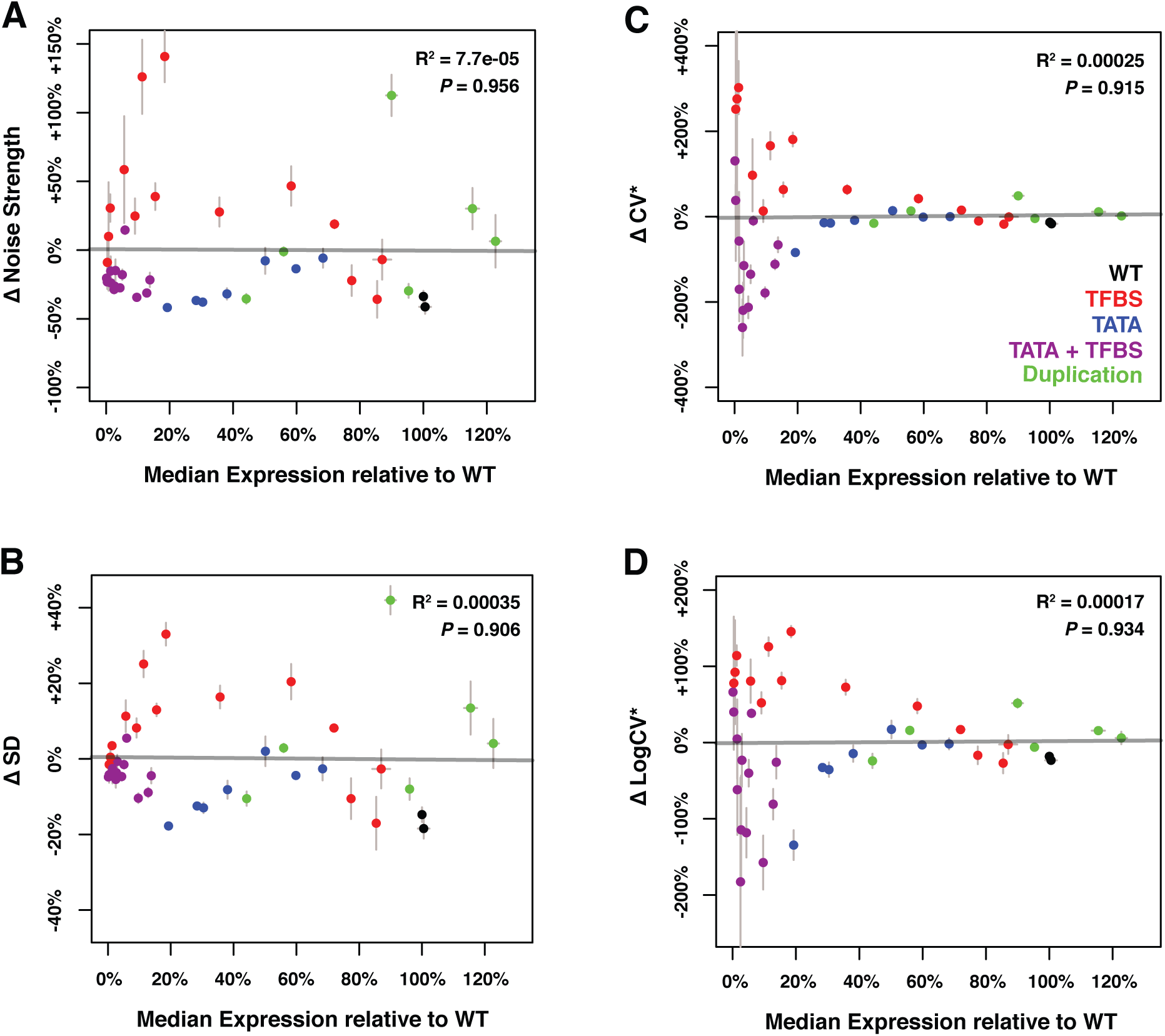
Relationship between median expression level and four different metrics of ΔNoise. ΔNoise was calculated as the residuals of a LOESS regression of expression noise on median expression, using four different metrics of noise: **(A)** Noise strength, equal to *SD*^2^/*median* **(B)** SD, the standard deviation of expression among genetically identical cells **(C)** CV*, equal to *SD/median* and **(D)**LogCV*, equal to *log*_2_(*SD/median)*. The best linear fit between median expression and ΔNoise is shown as a gray line, with the coefficient of determination (“R^2^”) and the significance of the Pearson’s correlation coefficient (“*P*”) indicated in the upper right. As expected, ΔNoise strength, ΔSD, ΔCV* and ΔLogCV* are all uncorrelated with median expression. Colors represent different types of mutations in the *TDH3* promoter, as indicated in the lower right of panel **(B)**. Error bars show 95% confidence intervals calculated from 4-6 replicates.

**Figure 3 – figure supplement 3.**
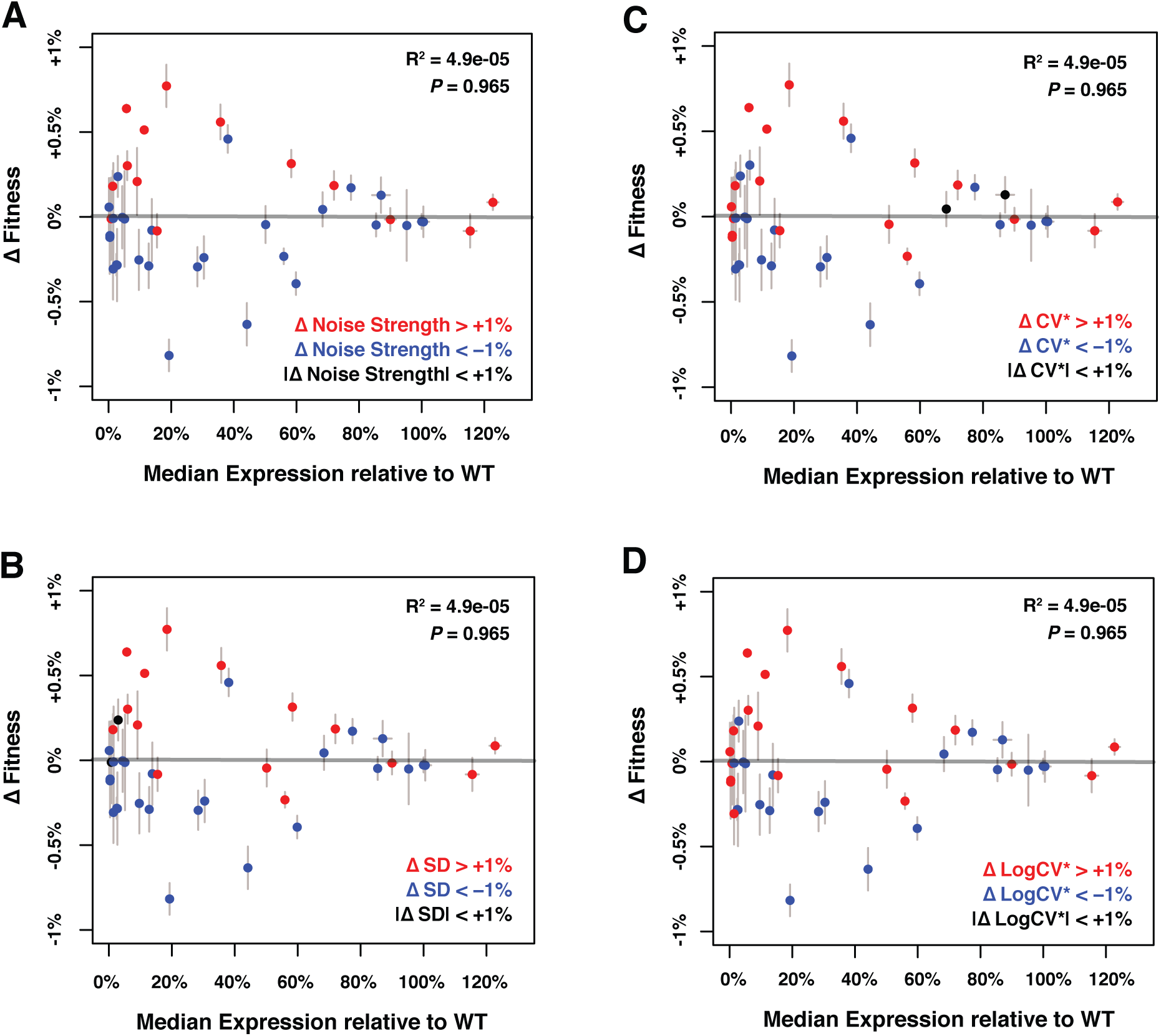
Relationship between median expression level and Δ Fitness. ΔFitness was calculated as the residuals of a LOESS regression of fitness on median *TDH3* expression. Genotypes were colored based on their level of noise measured as **(A)** Noise strength, equal to *SD*^2^/*median* **(B)** SD, the standard deviation of expression among genetically identical cells **(C)** CV*, equal to *SD/median* and **(D)** LogCV*, equal to *log*_2_(*SD/median*). The best linear fit between median expression and ΔFitness is shown as a gray line, with the coefficient of determination (“R^2^”) and the significance of the Pearson’s correlation coefficient (“*P*”) indicated in the upper right. As expected, ΔFitness is uncorrelated with median expression of *TDH3*. Error bars show 95% confidence intervals calculated from 4-6 replicates.

**Figure 3 – figure supplement 4.**
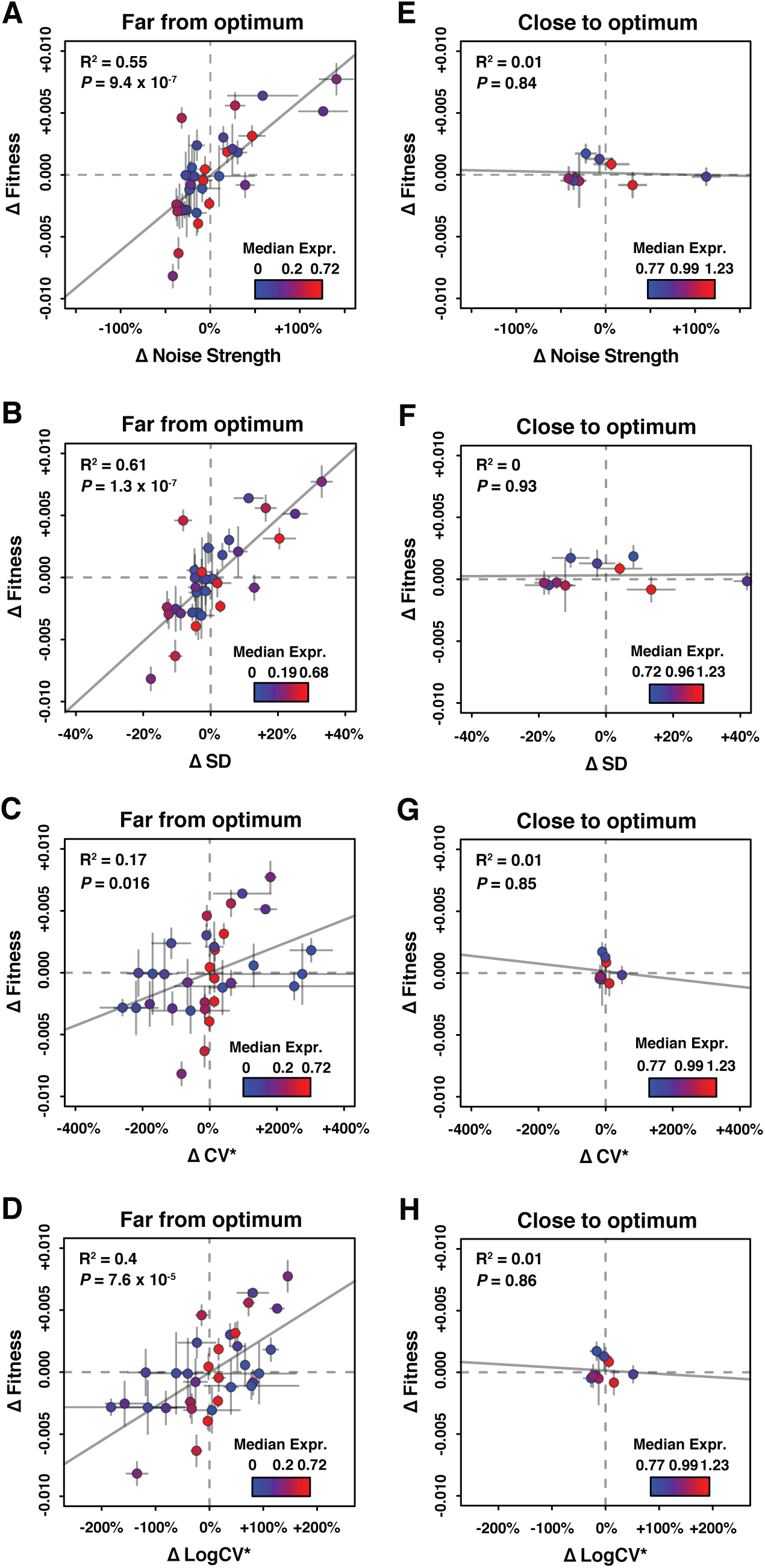
Relationship between ΔNoise and ΔFitness using four different metrics of noise. ΔNoise was calculated as the residuals of a LOESS regression of expression noise on median expression, using four different metrics of noise: **(A,E)** Noise strength, equal to *SD*^2^/*median* **(B,F)** SD, the standard deviation of expression among genetically identical cells **(C,G)** CV*, equal to *SD/median* and **(D,H)** LogCV*, equal to *log*_2_(*SD*/*median*). **(A-D)** Relationship between ΔNoise and ΔFitness when median expression is far from optimum. **(E-H)** Relationship between ΔNoise and ΔFitness when median expression is close to optimum. **(A-H)** The best linear fit between ΔNoise and ΔFitness is shown as a gray line, with the coefficient of determination (“R^2^”) and the significance of the Pearson’s correlation coefficient (“*P*”) indicated in the upper left of each panel. Dots are colored based on median expression levels of the corresponding *P*_*TDH3*_ alleles as indicated by color gradient. Error bars show 95% confidence intervals calculated from at least four replicate populations.

**Figure 3 – figure supplement 5.**
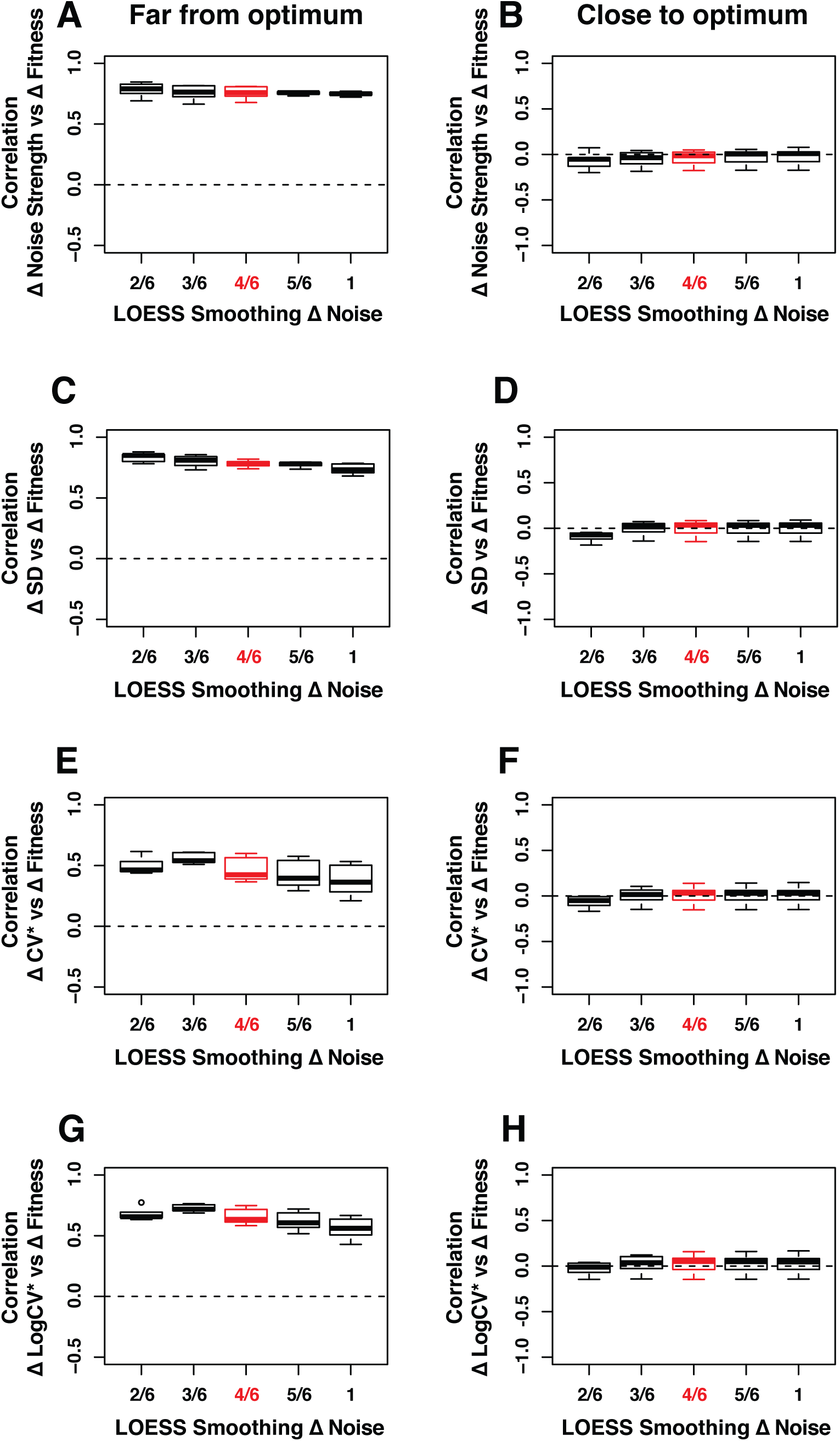
Robustness of the correlation between ΔNoise and ΔFitness to variation in the smoothing parameter of the LOESS regression used to compute ΔNoise. ΔNoise was calculated as the residuals of a LOESS regression of expression noise on median expression, using four different metrics of noise: **(A.B)** Noise strength, equal to *SD*^2^/*median* **(C,D)** SD, the standard deviation of expression among genetically identical cells **(E,F)** CV*, equal to *SD/median* and **(G,H)** LogCV*, equal to *log*_2_(*SD/median)*. The Pearson’s Correlation Coefficient (PCC) between ΔNoise and ΔFitness was computed for 100 combinations of three parameters for **(A,C,E,G)** strains with a *TDH3* expression level far from optimum and **(B,D,F,H)** strains with a *TDH3* expression level close to optimum. The three parameters that were varied were (1) the smoothing parameter of the LOESS regression used to compute ΔNoise (2/6, 3/6, 4/6, 5/6 and 1), (2) the smoothing parameter of the LOESS regression used to compute ΔFitness (2/6, 3/6, 4/6, 5/6 and 1) and (3) the fitness threshold used to classify strains as far from optimum or close to optimum (0.0025, 0.005, 0.075, 0.01). For each plot, the five boxes represent the variation of PCC between ΔNoise and ΔFitness when parameter 1 is fixed to one of the values shown on the x-axis and the two other parameters are allowed to vary. The thick horizontal lines represent the median PCC across the 20 combinations of parameters 2 and 3. The bottom and top lines of each box represent the 25^th^ and 75^th^ percentiles. Parameter values used in the main figures are shown in red.

**Figure 3 – figure supplement 6.**
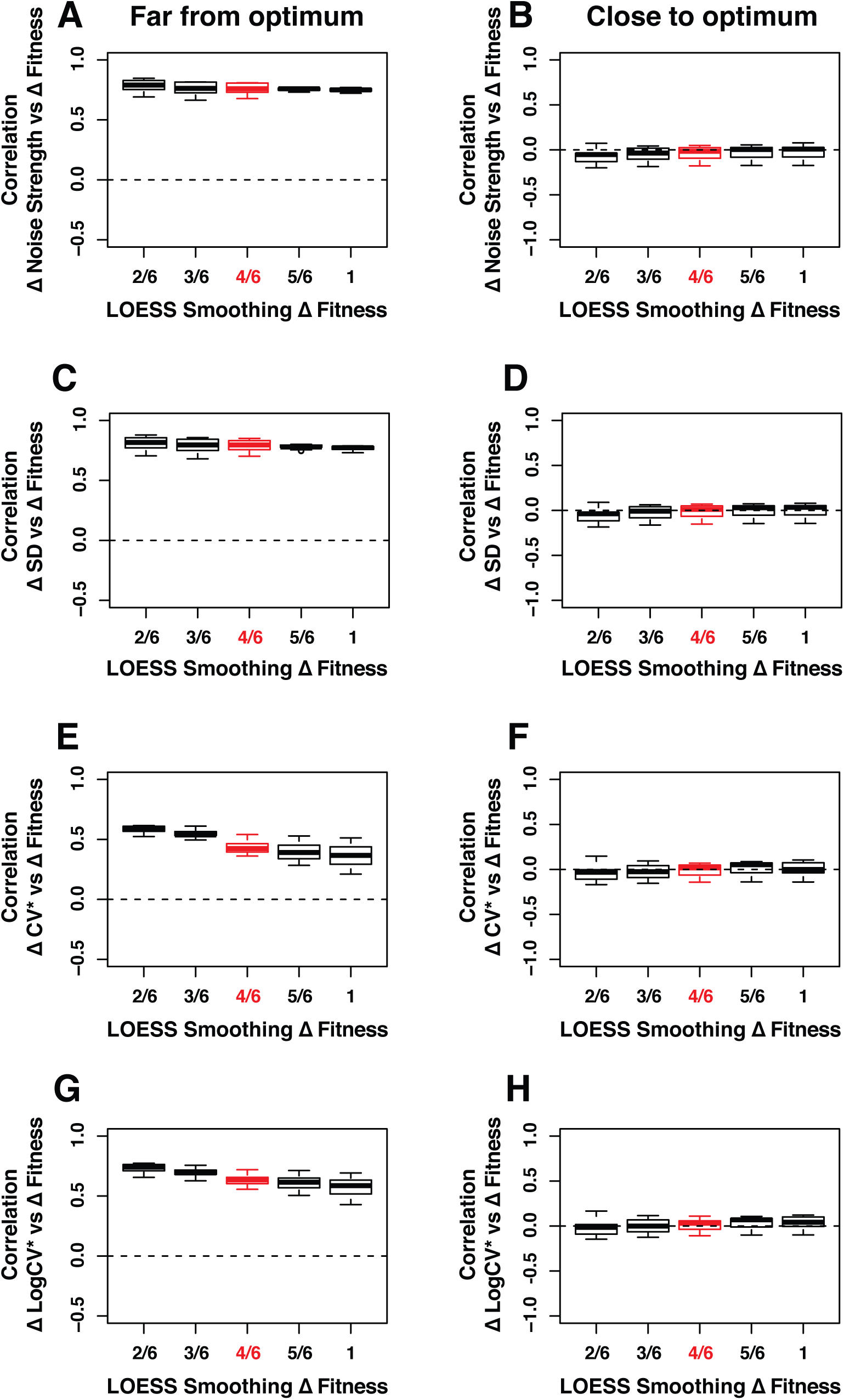
Robustness of the correlation between ΔNoise and ΔFitness to variation in the smoothing parameter of the LOESS regression used to compute ΔFitness. ΔNoise was calculated as the residuals of a LOESS regression of expression noise on median expression, using four different metrics of noise: **(A,B)** Noise strength, equal to *SD^2^/median* **(C,D)** SD, the standard deviation of expression among genetically identical cells **(E,F)** CV*, equal to *SD/median* and **(G,H)** LogCV*, equal to *log_2_(SD/median)*. The Pearson’s Correlation Coefficient (PCC) between ΔNoise and ΔFitness was computed for 100 combinations of three parameters for **(A,C,E,G)** strains with a *TDH3* expression level far from optimum and **(B,D,F,H)** strains with a *TDH3*expression level close to optimum. The three parameters that were varied were (1) the smoothing parameter of the LOESS regression used to compute ΔNoise (2/6, 3/6, 4/6, 5/6 and 1), (2) the smoothing parameter of the LOESS regression used to compute ΔFitness (2/6, 3/6, 4/6, 5/6 and 1) and (3) the fitness threshold used to classify strains as far from optimum or close to optimum (0.0025, 0.005, 0.075, 0.01). For each plot, the five boxes represent the variation of PCC between ΔNoise and ΔFitness when parameter 2 is fixed to one of the values shown on the x-axis and the two other parameters are allowed to vary. The thick horizontal lines represent the median PCC across the 20 combinations of parameters 1 and 3. The bottom and top lines of each box represent the 25^th^ and 75^th^ percentiles. Parameter values used in the main figures are shown in red.

**Figure 3 – figure supplement 7.**
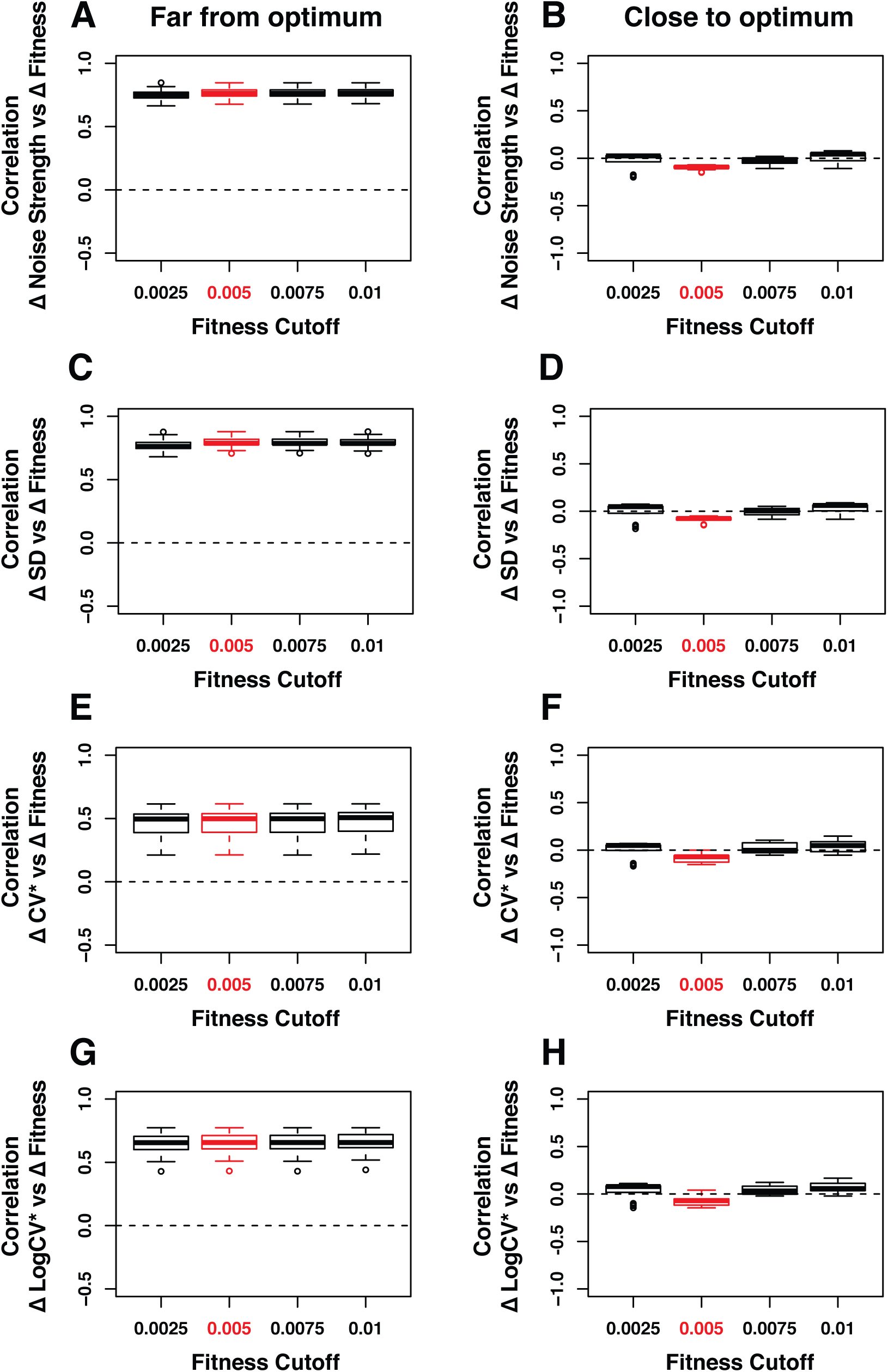
Robustness of the correlation between ΔNoise and ΔFitness to variation in the fitness threshold used to classify genotypes as far or close to optimum. ΔNoise was calculated as the residuals of a LOESS regression of expression noise on median expression, using four different metrics of noise: **(A,B)** Noise strength, equal to *SD^2^/median* **(C,D)** SD, the standard deviation of expression among genetically identical cells **(E,F)** CV*, equal to *SD/median* and **(G,H)** LogCV*, equal to *log*_2_(SD/median). The Pearson’s Correlation Coefficient (PCC) between ΔNoise and ΔFitness was computed for 100 combinations of three parameters for **(A,C,E,G)** strains with a *TDH3* expression level far from optimum and **(B,D,F,H)** strains with a *TDH3* expression level close to optimum. The three parameters that were varied were (1) the smoothing parameter of the LOESS regression used to compute ΔNoise (2/6, 3/6, 4/6, 5/6 and 1), (2) the smoothing parameter of the LOESS regression used to compute ΔFitness (2/6, 3/6, 4/6, 5/6 and 1) and (3) the fitness threshold used to classify strains as far from optimum or close to optimum (0.0025, 0.005, 0.075, 0.01). For each plot, the five boxes represent the variation of PCC between ΔNoise and ΔFitness when parameter 3 is fixed to one of the values shown on the x-axis and the two other parameters are allowed to vary. The thick horizontal lines represent the median PCC across the 25 combinations of parameters 1 and 2. The bottom and top lines of each box represent the 25^th^ and 75^th^ percentiles. Parameter values used in the main figures are shown in red.

**Figure 3 – figure supplement 8.**
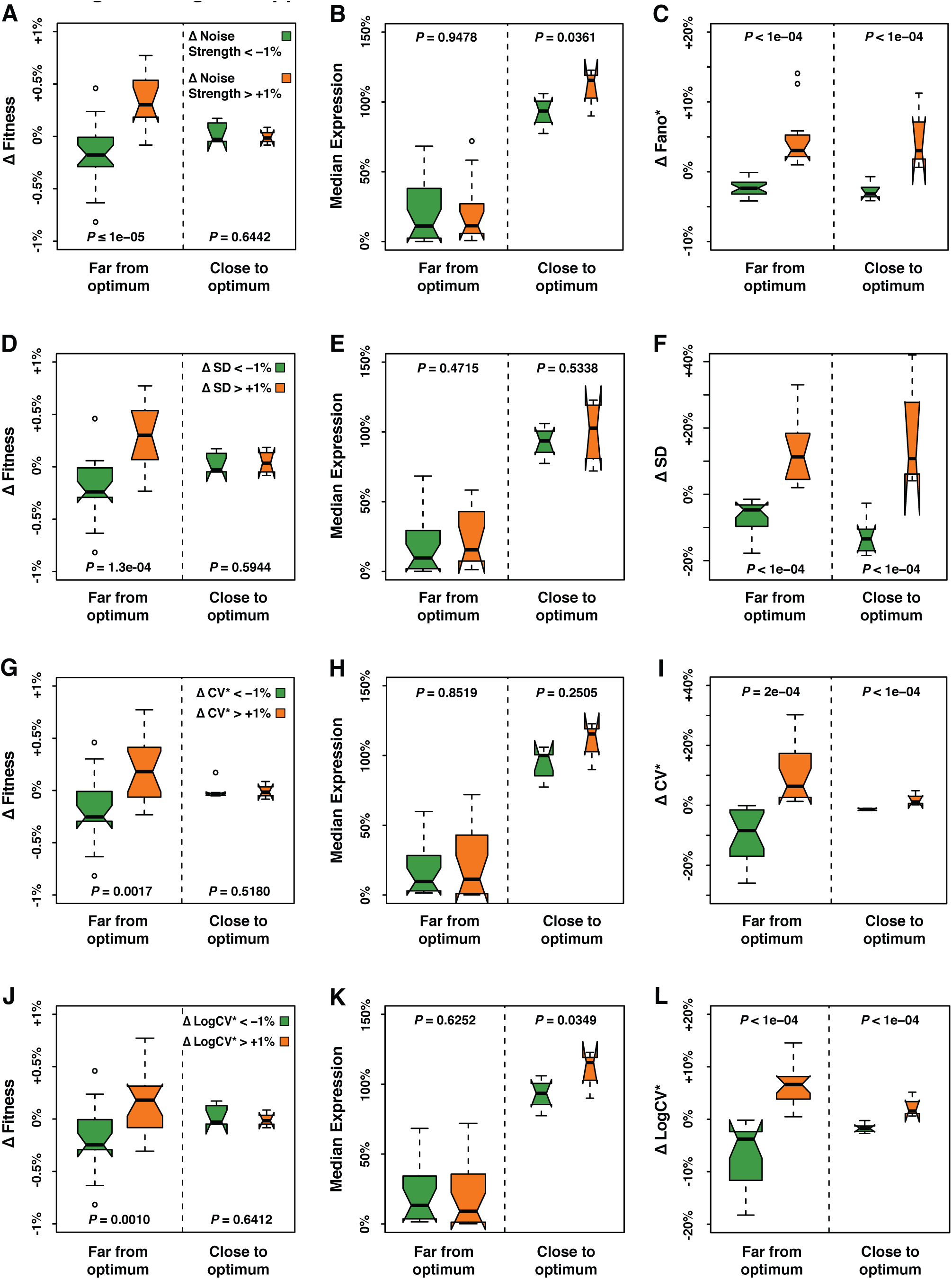
Fitness, median expression and noise of genotypes with ΔNoise above +1% compared to genotypes with ΔNoise below −1%. ΔNoise was calculated as the residuals of a LOESS regression of expression noise on median expression using four different metrics of noise: **(A-C)** Noise strength, equal to *SD^2^/median* **(D-F)** SD, the standard deviation of expression among genetically identical cells **(G-I)** CV*, equal to *SD/median* and **(J-L)** LogCV*, equal to *log_2_(SD/median)*. **(A,D,G,J)** Comparison of ΔFitness between genotypes with low ΔNoise (green) and genotypes with high ΔNoise (orange). ΔFitness was calculated as the residuals of a LOESS regression of fitness on median expression. **(B,E,H,K)** Comparison of median expression level between genotypes with low ΔNoise (green) and genotypes with high ΔNoise (orange). **(C,F,I,L)** Comparison of ΔNoise between genotypes with low ΔNoise (green) and genotypes with high ΔNoise (orange). **(A-L)** All comparisons were performed separately for genotypes with a median expression level close to optimum and genotypes with a median expression level far from optimum. ΔFitness, ΔNoise and the distance to the optimum expression level were determined independently for each of the four metrics of noise, as shown in Figure 3 – figure supplement 1. Thick horizontal lines represent the median ΔFitness across genotypes and the notches display the 95% confidence interval of the median. The bottom and top lines of each box represent the 25^th^ and 75^th^ percentiles. Permutation tests were used to assess the significance of the difference in phenotypes (ΔFitness, median expression and ΔNoise) between genotypes with low ΔNoise and high ΔNoise. For each test, the values of ΔNoise were randomly shuffled among genotypes 100,000 times. The *P* values shown below each plot represent the proportion of permutations for which the difference in median phenotype between genotypes with low and high ΔNoise was greater than the observed difference in median phenotype between genotypes with low and high ΔNoise.

**Figure 4 – figure supplement 1.**
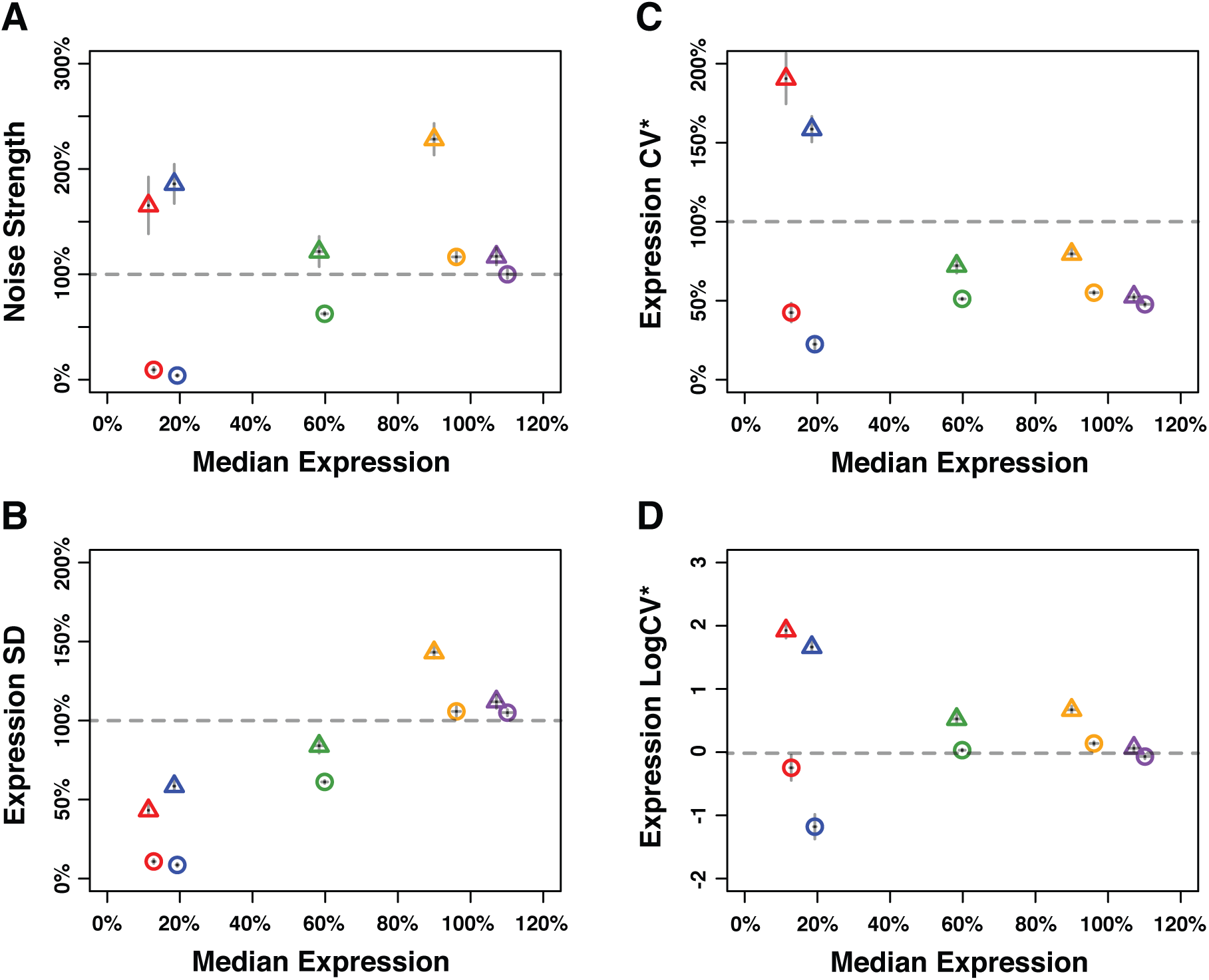
Median expression level and expression noise for five pairs of genotypes that were competed directly against each other. Four metrics of noise were used: **(A)** the variance of expression divided by the median expression (noise strength), **(B)** the standard deviation of expression among genetically identical cells (SD), **(B)** the standard deviation divided by the median expression (CV*) and **(D)** the binary logarithm of the standard deviation divided by the median expression (LogCV*). **(A-D)** Different colors are used to distinguish pairs of genotypes (*P*_*TDH3*_ alleles) with similar median expression levels. Each pair comprises one genotype with low expression noise (circle) and one genotype with high expression noise (triangle). Error bars show 95% confidence interval obtained from at least 3 replicates.

**Figure 6 – figure supplement 1.**
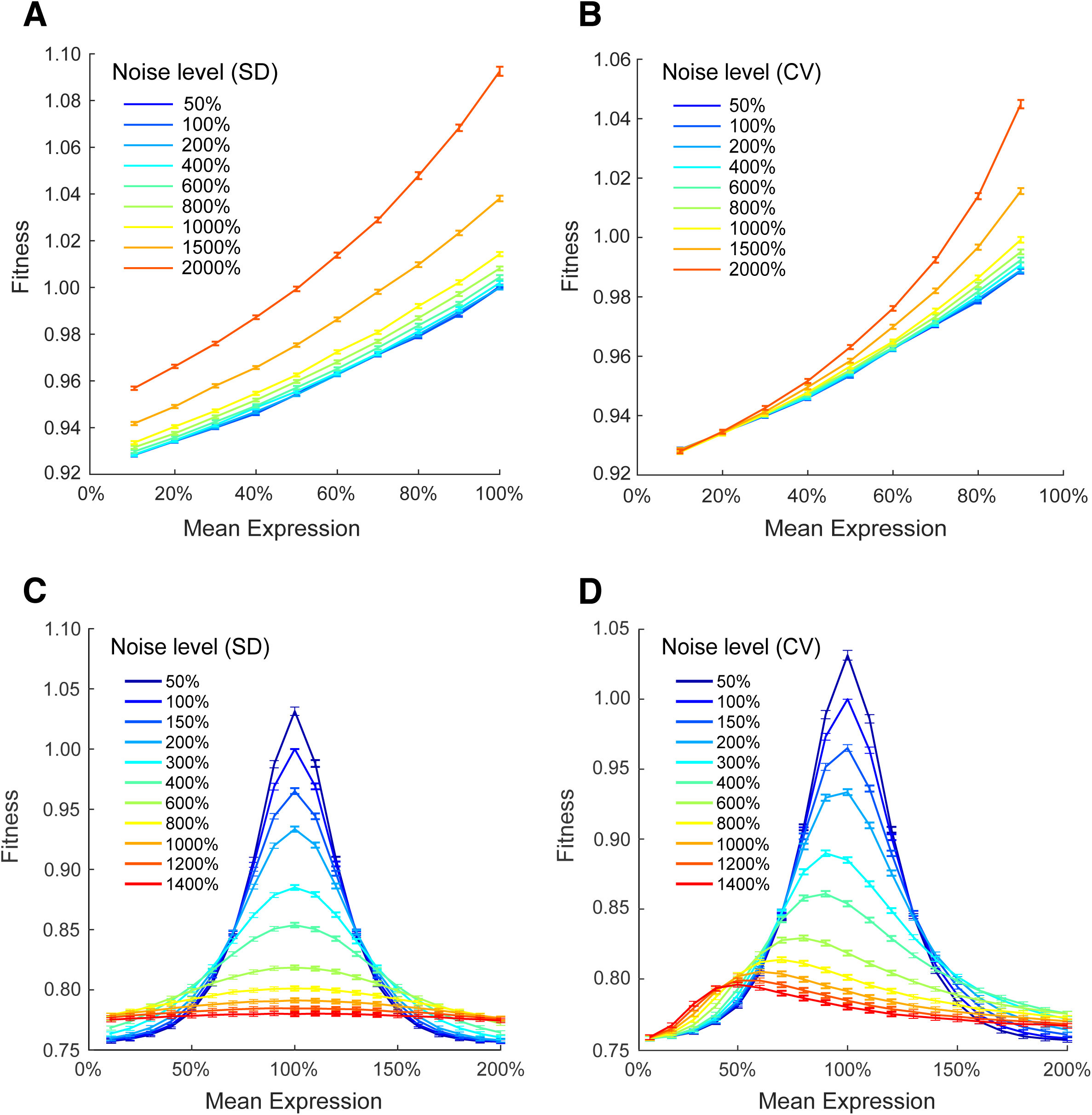
Simulating the effect of two different metrics of expression noise on fitness at different median expression levels. Population fitness was simulated for median expression levels *μ*_*E*_ ranging from 10% to 100% and for: **(A)** the standard deviation of expression *σ*_*E*_ ranging from 0.05 to 2 and the linear function *DT* = −40 × *E* + 160 relating single cell expression to doubling time, **(B)** the coefficient of variation of expression *σ*_*E*_/*μ*_*E*_ ranging from 0.05 to 2 and the linear function *DT* = −40 × *E* + 160 relating single cell expression to doubling time, **(C)** the standard deviation of expression *σ*_*E*_ ranging from 0.05 to 0.8 and the Gaussian function *DT* = −160 × exp(− (*E* − 1)^2^/0.18) + 240 relating single cell expression to doubling time, and **(D)** the coefficient of variation of expression *σ*_*E*_/*μ*_*E*_ ranging from 0.05 to 1.4 and the Gaussian function *DT* = −160 × exp(−(*E* − 1)^2^/0.18) + 240 relating single cell expression to doubling time. **(A-D)** Error bars show 95% confidence intervals of mean fitness calculated from 100 replicate simulations for each combination of mean expression and expression noise.

**Figure 6 – figure supplement 2.**
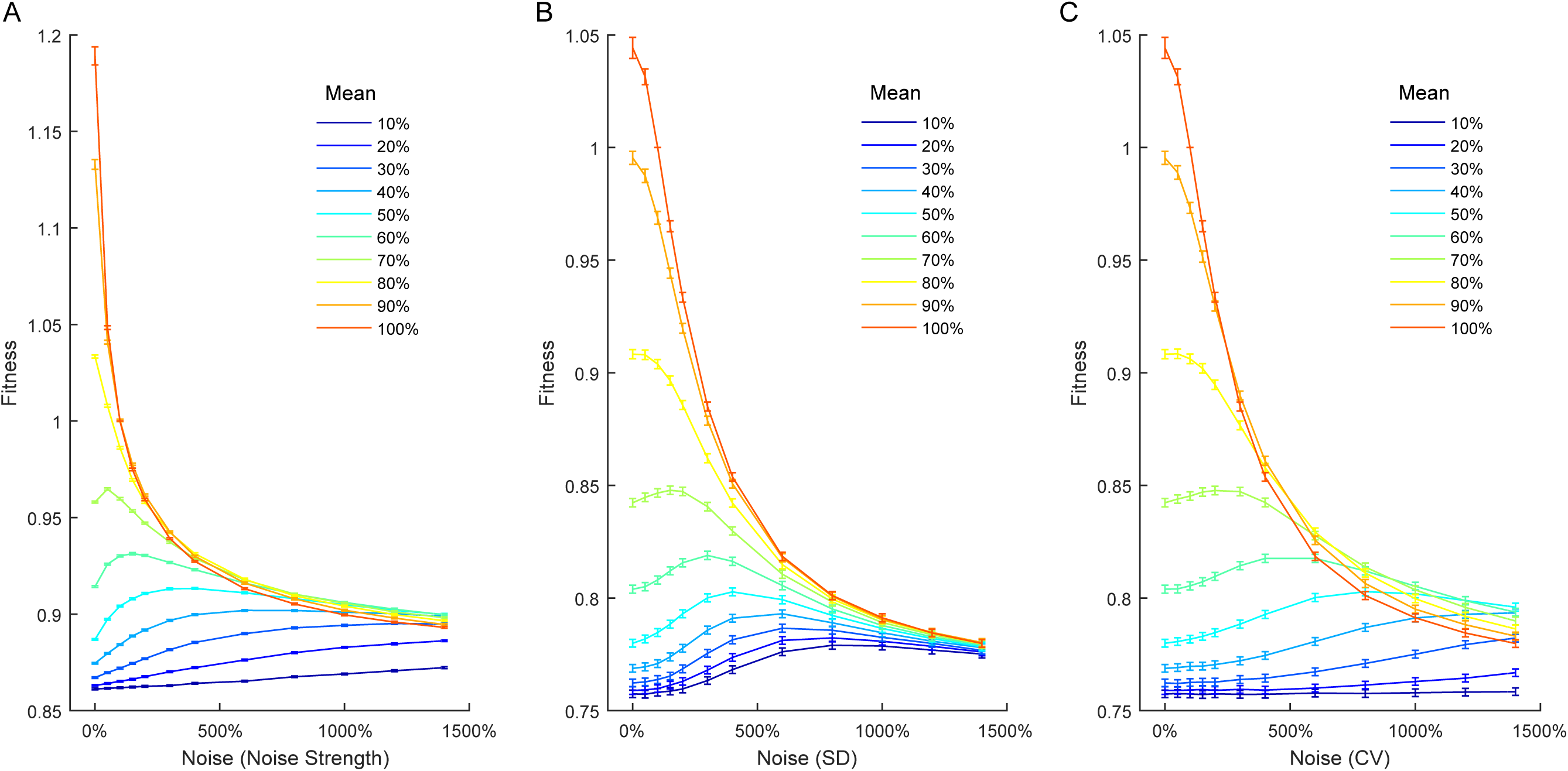
Relationship between expression noise and fitness at different values of mean expression in simulations using a Gaussian function relating single cell expression to doubling time. Three different noise metrics were used: **(A)** the noise strength *σ*_*E*_^2^/*μ*_*E*_, **(B)** the standard deviation *σ*_*E*_, **(C)** the coefficient of variation *σ*_*E*_/*μ*_*E*_. **(A-C)** Error bars show 95% confidence intervals of mean fitness calculated from 100 replicate simulations for each combination of mean expression and expression noise.

